# PPARγ controls ESCRT-dependent fibroblast-like synoviocyte exosome biogenesis and Alleviates Chondrocyte Osteoarthritis Mediated by Exosomal ANXA1

**DOI:** 10.1101/2023.11.19.567728

**Authors:** Shuangshuo Jia, Yue Yang, Jiabao Liu, Yingliang Wei, Ziyuan Wang, Lingyu Yue, Lunhao Bai

## Abstract

Osteoarthritis (OA) is characterized by synovitis, cartilage degeneration and exercise therapy has been recognized as first line therapy. The exercise related exosome involved in the interaction between fibroblast-like synoviocytes (FLSs) and chondrocytes could be a novel promising strategy for treating OA. In this study, PPARγ was upregulated in FLSs under exercise by single cell transcriptome sequencing. Then, we investigated the underlying mechanisms of PPARγ-treated FLSs derived exosome on OA in vivo and vitro. Our data revealed that overexpression PPARγ FLSs derived exosome could ameliorate the OA severity in vivo and vitro. But knockdown PPARγ FLSs derived exosome aggravate OA. Moreover, we found PPARγ controls the endosomal sorting complex required for the transport (ESCRT)-dependent exosome biogenesis pathway. Annexin A1 (ANXA1) was enriched in OE-PPARγ exosome by quantitative proteomics. By Chip-qPCR and Co-IP methods, PPARγ and its coactivator -1α (PGC-1α) acts with ESCRT subunits including HRS, STAM1, TSG101, CHMP7 and promotes their association to cargo ANXA1. As a therapeutic cargo, exosomal ANXA1 was confirmed be internalization by chondrocyte via exosome labeled experiment and ANXA1 could inhibit the phospharylation of ERK to activate the autophagy and decrease chondrocyte apoptosis. While the ANXA1 receptor blocker BOC-2 could reverse the therapic effect. In conclusion, PPARγ/ESCRT – FLSs exosomal ANXA1 – ERK axis provides a deeper theoretical basis for exercise therapy of OA and a new idea for the clinical transformation of exosomes into OA therapy.

## 1. Introduction

Osteoarthritis (OA) is a chronic degenerative joint disorder that results in disability and impacts more than 500 million individuals globally [1, 2]. It is typified by cartilage degradation and synovitis, which are directly linked to clinical symptoms like joint swelling and inflammatory pain [3–5]. The synovium houses metabolically highly active cells known as fibroblast-like synoviocytes (FLSs). These cells contribute to the unique functional properties of articular surfaces, provide nutrients to the cartilage, constitute the chondrocytes’ microenvironment, and regulate chondrocyte activity [6, 7]. Therefore, understanding the intercellular communication between FLSs and chondrocytes during OA development may illuminate new therapeutic strategies.

Current clinical guidelines widely recommend non-surgical therapies like physical exercise as first-line treatments [8]. Our prior research has established a moderate treadmill exercise protocol and demonstrated its beneficial effects in ameliorating OA [9–13]. We further identified the potential role of peroxisome proliferator-activated receptor γ (PPARγ), a common upstream element in our sequential studies on irisin [9] and Meteorin-like (metrnl) [10] in OA exercise therapy. Exercise induces PPARγ and its coactivator-1α (PGC1α), stimulating many well-known beneficial effects of exercise [14]. Although PPARγ’s potential involvement in OA has been reported [15], the mechanism by which its upregulation in FLSs impacts articular chondrocytes during exercise therapy for OA remains unclear.

FLSs release exosomes [16, 17]. Exosomes are extracellular vesicles (EVs) approximately 30 to 200 nm in diameter and with a density between 1.13 and 1.19 g/mL. They mirror the cell’s topology and are enriched in select proteins, lipids, nucleic acids, and glycoconjugates [18, 19]. These exosomes serve as crucial mediators of intercellular communication and participate in many physiological and pathological processes [20]. Nevertheless, the impact of exosomes mediated by exercise-related genes (PPARγ) on articular chondrocytes remains unexplored. Also, the regulation mechanisms that determine exosome contents during exosome biogenesis mediated by PPARγ remain poorly understood. Further, bioactive cargoes in exosomes can be uptaken by recipient cells, altering their physiological and pathological states [18]. Thus, it is worthwhile to explore which cargoes in exosomes play a role in OA and how PPARγ participates in FLSs related exosome biogenesis.

Exosome biogenesis is a complex multistep process including the formation of early endosomes through endocytosis of the cytoplasmic membrane, late endosomal multivesicular bodies (MVBs) formation by intraluminal vesicles (ILVs) bud inward from endosomal compartments, and MVBs fuse to the cell surface and release their ILVs as exosomes [21, 22]. During the maturation of endosomes from early to late stages, endosomal proteins and macromolecules are selectively sorted in the ILVs of MVBs. The protein complex called endosomal sorting complex required for transport (ESCRT) is crucial in generating these unique membrane compartments [21, 23]. ESCRTs consists of four multimeric protein complexes (ESCRT-0, -I, -II, and -III) which play a key role in protein sorting and also important in ILV biogenesis and MVB formation for exosome biogenesis [24].

Annexin 1 (ANXA1), a 37 kDa member of the annexin superfamily of Ca2+ dependent phospholipid-binding proteins, is linked to membrane-related events and cellular functions [25]. As an endogenous mediator, ANXA1 plays an important role in resolving inflammation, autophagy, and apoptosis [26]. ANXA1 has been found in exosomes and is involved in intercellular communication [27–29]. Since ANXA1 expression is positively correlated with PPARγ, and PPARγ ligands may induce annexin A1 expression [30], we hypothesized that PPARγ overexpression in FLS-derived exosomes may impact OA via exosomal ANXA1.

During OA, cytokines activate the extracellular signal-regulated kinase (ERK), a serine/threonine protein kinase [31]. ERK primarily regulates apoptosis and autophagy [32]. Inhibition of ERK triggers an upregulation of autophagic processes [33]. Autophagy, a cellular self-protection mechanism, serves as an intracellular scavenger to maintain cellular homeostasis [34, 35] and plays a significant role in the pathological development of OA [34]. Chondrocyte apoptosis is implicated in cartilage degeneration pathogenesis in osteoarthritis, with autophagy facilitating the removal of senescent cells in aging cartilage [36, 37].

In this study, we identified an unknown function of PPARγ in exosome biogenesis and the mechanisms for the regulation of ESCRT-dependent exosome formation. We also assess the mechanism of exercise-induced FLSs PPARγ-exosomal ANXA1-ERK axis in managing apoptosis, anabolism, and catabolism of OA via autophagy activation.

## 2. Results

### 2.1 PPARγ was Up-regulated in the Synovium of OA-affected SD rats After Exercise Therapy

The synovium, cartilage, and subchondral bone were analyzed using single-cell transcriptome sequencing. The t-distributed stochastic neighbor embedding (t-SNE) projection (Supplementary figure 1A) identified 11 distinct cell clusters, including chondrocytes, FLSs, B cells, macrophages, NK cells, among others. This study, however, primarily focused on the differential gene expression in FLSs. We isolated 5213 (29.83%) FLSs in the control group (CG), 9513 (47.49%) FLSs in the OA group, and 5633 (33.13%) FLSs in the exercise and OA group (EXE+OA). In the EXE+OA group, 211 genes were downregulated and 247 genes were upregulated compared to the OA group. The t-SNE projection (Supplementary figure 1B), heatmap cluster analyses (Supplementary figure 1C), violin plot (Supplementary figure 1D), volcano plots (Supplementary figure 1E), and highlighted the top 9 differentially expressed genes (DEGs) between the EXE+OA and OA groups. PPARγ was found to be upregulated in the EXE+OA group.

As illustrated in Supplementary figure 2B and 2C, when comparing the CG with the OA groups, 480 genes were downregulated and 509 genes were upregulated in the OA group. Among all upregulated genes, PPARγ also exhibited an increase in the CG group, albeit not as pronounced as in the EXE+OA group. Collectively, these results underscore the significance of PPARγ at the single-cell transcriptome level during exercise therapy for OA.

### 2.2 PPARγ was Downregulated in the synovium of patients with increasing Kellgren-Lawrence grade and was related to Pathological Features

In order to assess the potential role of PPARγ in OA, we initially confirmed the integrity of articular synovium specimens through imaging evaluation, histological evaluation and OA-related protein analysis (Figure 1). By integrating western blot analyses (Figure 1C, D) with histological evaluations (Figure 1A), we discerned that PPARγ was notably downregulated (p < 0.001) within the synovium of increasing Kellgren-Lawrence (K-L) grade which indicates more severer OA and less PPARγ.

**Figure 1.**
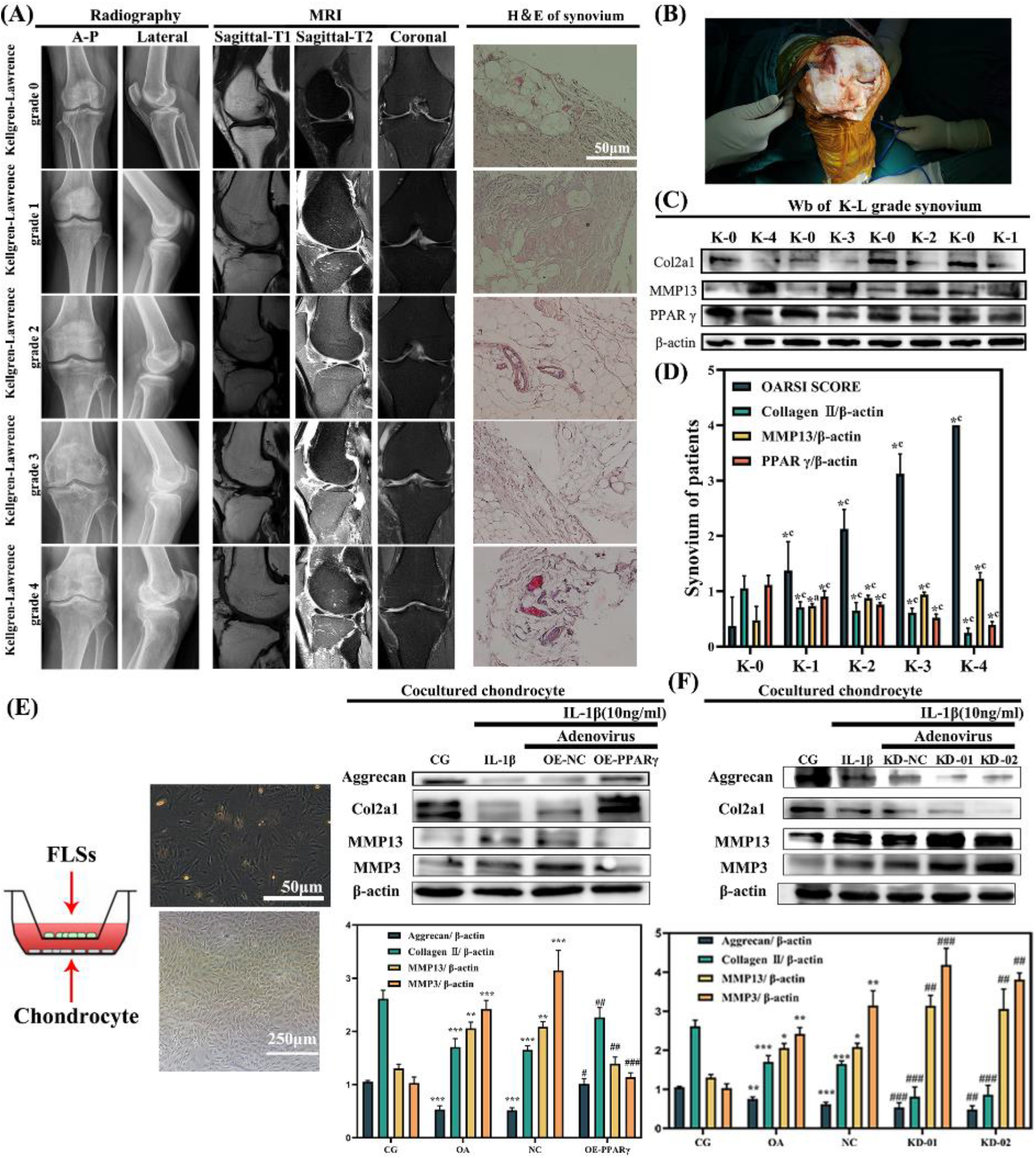
Expression of PPARγ is downregulated in synovium with increasing Kellgren-Lawrence grade. (A) Plain radiograph, magnetic resonance imaging (MRI) scans and hematoxylin & eosin (H&E). A–P: anterior–posterior. (B) Total knee arthroplasty. (C) Western blots of PPARγ in synovial tissue. (D) Comparison of all Kellgren-Lawrence grades against grade 0 (*P < 0.001, *a p < 0.01, *b p < 0.01, *c p < 0.001; ANOVA) shown. (E) Co-culturing overexpressed or knocked-down (OE/KD) PPARγ-treated FLSs with primary chondrocytes underscored the significance of PPARγ and intercellular communication. (F) The ratio of phenotype changes in co-cultures of OE/KD PPARγ-treated FLSs with primary chondrocytes. Values are expressed as mean ± SEM. *p < 0.05 vs. CG; **p < 0.01 vs. CG; ***p < 0.001 vs. CG. #p < 0.05 vs. OA; ##p < 0.01 vs. OA; ###p < 0.001 vs. OA. Means ± 95 % CI. β-actin was used as an internal control. Data are presented as mean ± 95 % CI; n = three patients per group. K-0 refers to Kellgren-Lawrence grade 0.

### 2.3 Co-culture of OE/KD PPARγ-treated FLSs with Primary Chondrocytes Revealed the Importance of PPARγ and Intercellular Communication in OA Therapy

As depicted in Figure 1E, under co-culture conditions, FLSs treated with OE-PPARγ mitigated IL-1β-induced inflammation in chondrocytes, as evidenced by increased aggrecan and COL2α1, and decreased MMP13 and MMP3. Conversely, FLSs treated with KD-01 or KD-02 exacerbated the inflammation in co-cultured chondrocytes. Notably, the inflammatory effect of IL-1β in chondrocytes was counteracted by co-culturing with FLSs overexpressing PPARγ but was exacerbated by knockdown treatments. There were no significant differences between the OA+NC groups or the OA group. The results from the co-culture experiment sparked a keen interest in the intercommunication between FLSs and chondrocytes (Figure 1E and 1F). The overexpression and knockdown efficiency of PPARγ, both *in vivo* and *in vitro*, were validated using quantitative real-time PCR (qRT-PCR) (refer to Supplementary figure 2D).

### 2.4 Exosomes from PPARγ-treated FLSs and Evidence of Internalization by Chondrocytes

After co-culturing chondrocytes with OE/KD PPARγ-treated FLSs, we examined whether the exosomes secreted by FLSs play a critical role in their interaction with chondrocytes. TEM revealed a hollow, sphere-like morphology of OE/KD PPARγ-treated FLS-derived exosomes (Figure 2A). We utilized NTA to assess the size distribution of these exosomes (Figure 2B). The primary peak in particle size was observed between 60 and 130 nm, with 92.0% of the overall size distribution ranging from 30 to 200 nm. The average size was determined to be 136.5±2.2 nm, and the concentration was approximately 6.14×10^9^ particles/mL (Figure 2B). To ascertain whether chondrocytes could internalize FLS-derived exosomes, we labeled these exosomes with PKH-67 (red) and PKH-26 (green). Confocal microscopy revealed that the PKH-67 and PKH-26 double-labeled exosomes were localized in the perinuclear areas of the chondrocytes (Figure 2G), thereby confirming their internalization by chondrocytes. Moreover, the protein expression of exosomal markers, namely ALIX, CD9, and CD63 (Figure 2H), was found to be enriched in the exosome groups, suggesting successful exosome isolation. The calnexin was used for exosome negative control.

**Figure 2.**
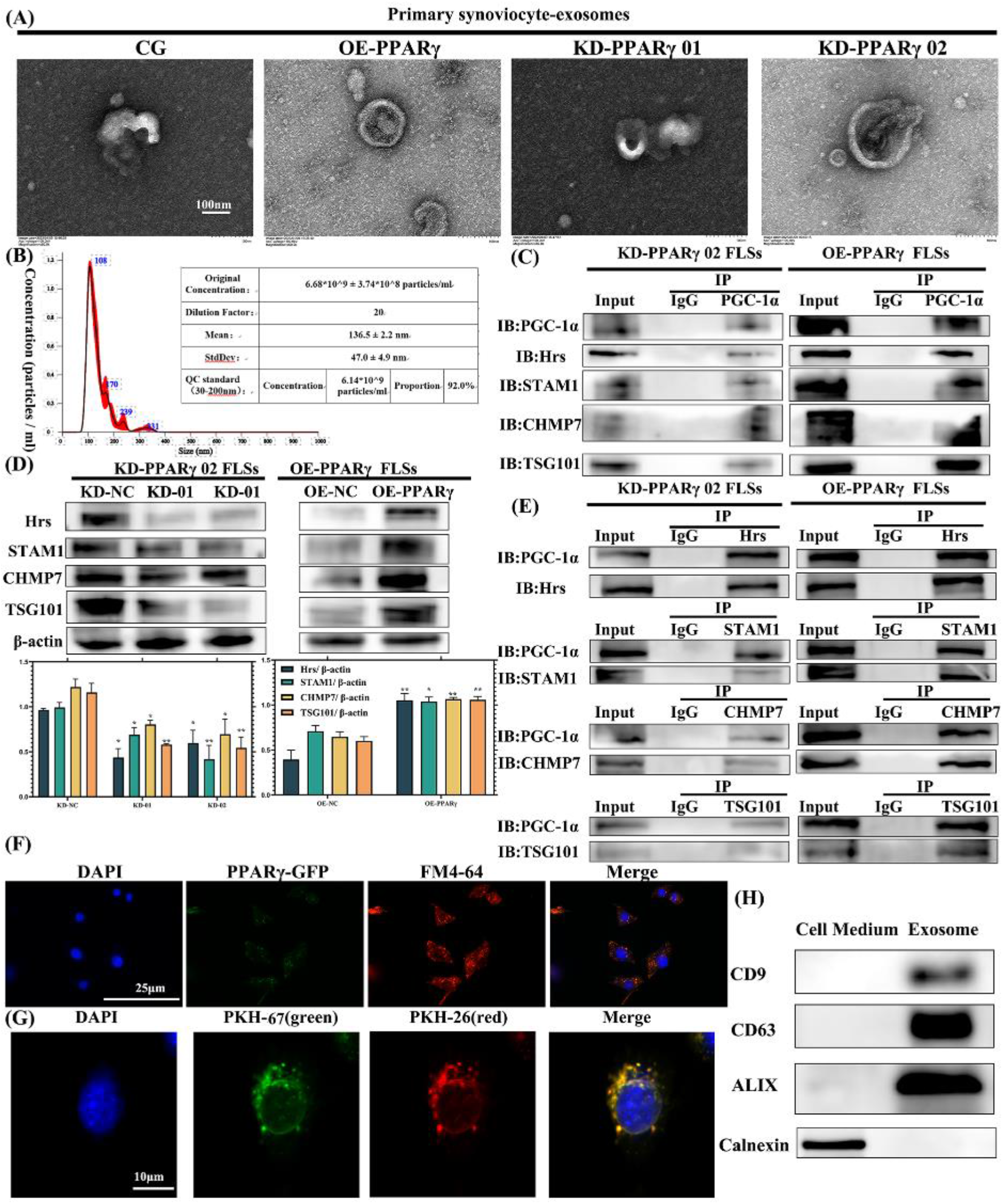
Isolation and characterization of exosomes derived from FLSs, PPARγ interacts with exosome biogenesis proteins and regulates exosome secretion through ESCRT-dependent pathway. (A) Transmission electron microscopy (TEM) image of exosomes derived from FLSs. (B) Nanoparticle tracking analysis (NTA) was used to assess the size distribution of FLS-derived exosomes. The primary peak in particle size ranged from 60 to 130 nm, and 92.0% of the overall size distribution fell between 30 and 200 nm. The mean particle size was 136.5±2.2 nm, and the concentration was 6.14×10^9^ particles/mL. (C) PGC-1α immunoprecipitated from FLSs with anti-PGC-1α and anti-ESCRT (Hrs, STAM1, CHMP7, TSG101) antibodies respectively, were analyzed by immunoblotting. (D) Lower levels of ESCRTs in PPARγ-KD cells, but higher levels in OE groups. (E) IB-PGC-1α was detected in the IP-Hrs, STAM1, TSG101, CHMP7 groups. (F) Representative confocal microscopy images showing colocalization of PPARγ-GFP with endosomal vesicle structures (FM4-64, red) in FLSs. (G) FLS-derived exosomes, labeled with PKH-67(red) and PKH-26 (green), were found to be taken up by chondrocytes through an exosome tracking technique. Confocal microscopy revealed that PKH-67 and PKH-26 labeled exosomes were located in the perinuclear areas of chondrocytes. (H) Exosome markers (CD9, CD63, ALIX) were detected through western blot analysis. Calnexin was used for negative control.

### 2.5 PPARγ interacts with exosome biogenesis proteins and regulates exosome secretion through ESCRT-dependent pathway

To study the roles of PPARγ in exosome biogenesis, we explored the interaction between the PPARγ coactivator PGC-1α and several ESCRT subunits using Co-IP in OE/KD PPARγ-treated FLSs. As shown in Figure 2C, the Hrs (ESCRT-0 subunit), STAM1 (ESCRT-0 subunit), TSG101 (ESCRT-I subunit) and CHMP7 (ESCRT-III subunit) protein was detected in the IP-PGC-1α, and no PGC-1α protein was observed in the band of IgG binding protein. Additionally, IB-PGC-1α was detected in the IP-Hrs, STAM1, TSG101, CHMP7 groups (Figure 2E). This confirmed the interaction between the PPARγ coactivator PGC-1α and ESCRT subunits, suggesting that PPARγ interacts with exosome biogenesis proteins.

To further study the functions of PPARγ in exosome biogenesis, we expressed PPARγ in FLSs and analyzed its localization. We observed that PPARγ-GFP formed a punctate subcellular pattern that colocalized with FM4-64 (Figure 2F), which stains the endosomal vesicle structures and indicated PPARγ might participated in endocytosis pathway. Consistently, to examine the role of PPARγ in exosome production, we generated OE/KD PPARγ cell lines. We observed lower levels of Hrs, STAM1, TSG101 and CHMP7 in PPARγ-KD cells, but higher levels in OE groups (Figure 2D). Taken together, we found PPARγ regulated exosome secretion through ESCRT-dependent pathway.

### 2.6 Exosomes Derived from OE-PPARγ-treated FLSs Relieved OA both *in vivo* and *in vitro*, whereas Exosomes Derived from KD-PPARγ-FLSs Aggravated OA

*In vivo*, we evaluated our SD rats using H&E, toluidine blue staining, and safranin O-fast green (SO-FG) for histological observations (Figure 3A, 4A) following intra-articular exosome injections. The osteoarthritis (OA) and OE-NC groups exhibited cartilage damage and hypocellularity relative to the control group (CG). However, the OE-PPARγ group displayed a smoother and relatively intact cartilage surface in comparison to the OA and OE-NC groups, suggesting that OE-PPARγ exosome could alleviate OA symptoms. OARSI score-based histological analysis indicated that tibiofemoral joints in the OA and OE-NC groups had diminished and damaged cartilage compared to the CG. Yet, the articular cartilage damage was found to be reversed in the OE-PPARγ group compared to OA and OE-NC. The KD-01 and KD-02 groups, however, displayed more severe cartilage surface damage and higher OARSI scores than the OA and KD-NC groups.

**Figure 3.**
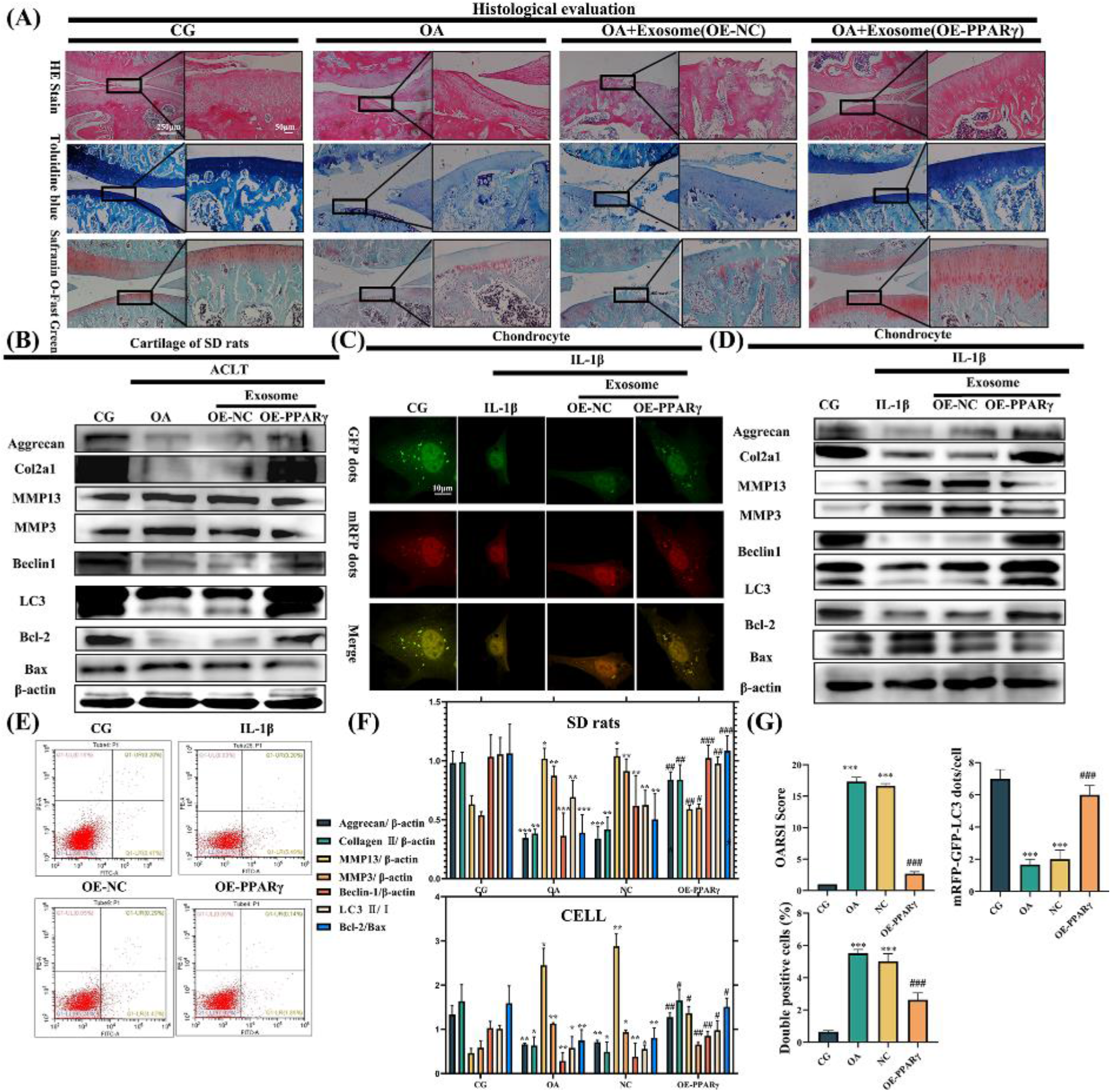
Exosomes derived from OE-PPARγ-treated FLSs can alleviate OA both *in vivo* and *in vitro*. (A) Histological analysis was performed using hematoxylin & eosin (H&E), toluidine blue, and safranin O-fast green staining. Higher magnification areas are represented by black boxes. We noted increased surface roughening, fibrillation, fissures, and erosions extending to the subchondral bone in the OA and OA+Exosome (OE-NC) groups compared with the control group (CG). A relatively smoother and more complete articular cartilage surface was observed in the OA+exosome (OE-PPARγ) group in comparison to the OA and OE-NC groups. (B) Relative protein expressions were assessed in the cartilage of SD rats and in chondrocytes (D) treated with OE-NC and OE-PPARγ exosomes. Aggrecan and COL2α1, which decrease in OA pathology, were restored by OE-PPARγ exosome treatment. Matrix metalloproteinase 13 (MMP13) and 3 (MMP3) decreased in the OE-PPARγ group compared to the OA group. Treatment with OE-PPARγ exosomes increased Beclin-1 and the LC3Ⅱ/Ⅰ ratio, indicating autophagy activation. The B-cell lymphoma-2 (Bcl-2) to its associated X protein (Bax) ratio was upregulated in the OE-PPARγ group, suggesting reduced chondrocyte apoptosis. (C) Chondrocytes treated with OE-PPARγ displayed mRFP-GFP-LC3 adenovirus double labeling. mRFP (red) indicates autolysosomes (ALs), while merged signals (yellow) represent autophagosomes (APs). (E) Flow cytometry analysis reveals reduced chondrocyte apoptosis in the OE-PPARγ group. (F) Displays the ratio of relative protein expression in SD rats and chondrocytes. (G) The OARSI score, the quantification of GFP and mRFP dots per cell in chondrocytes, and the quantification of flow cytometry analysis are presented. *p < 0.05, **p < 0.01, ***p < 0.001 vs. CG. #p < 0.05, ##p < 0.01, ###p < 0.001 vs. OA. Means ± 95 % CI.

We further evaluated the relative inflammatory proteins in the articular cartilage via immunohistochemistry (Figure 5A) and western blot analysis, both *in vivo* (Figure 3B, 4C) and *in vitro* (Figure 3D, 4B). The cartilage-specific proteins aggrecan and Col2α1, reduced by OA pathology, were restored by OE-PPARγ exosomes but worsened by KD-PPARγ exosomes. Inflammatory proteins, such as matrix metalloproteinase 13 (MMP13) and 3 (MMP3), were elevated in the KD-PPARγ exosome-treated group but diminished in the OE-PPARγ group compared to the OA group. As for autophagy and apoptosis-related indicators, Beclin-1 and microtubule-associated proteins light chain 3 (LC3) were utilized for autophagy level detection. Treatment with OE-PPARγ exosomes increased Beclin-1 and LC3Ⅱ/Ⅰ levels, signifying autophagy activation, whereas KD-PPARγ (KD-01, KD-02) suppressed this trend. The ratio of B-cell lymphoma-2 (Bcl-2) to its associated X protein (Bax) was upregulated in OE-PPARγ but downregulated in KD-PPARγ, indicating that OE-PPARγ could mitigate chondrocyte apoptosis, whereas KD-PPARγ would exacerbate apoptosis.

Furthermore, mRFP-GFP-LC3 adenovirus double labeling (Figure 3C and 4E) and transmission electron microscopy (TEM) (Figure 5C) revealed that autophagosomes (APs), characterized by double-layer or multilayer membranes encapsulating portions of the cytoplasm and organelles slated for degradation [38], were increased by OE-PPARγ exosomes, but decreased by KD-PPARγ exosomes. Flow cytometry results showed that apoptosis decreased following intervention with OE-PPARγ exosomes, but increased with KD-PPARγ exosomes (Figure 3E and 4F). Taken together, these findings suggest that exosomes derived from PPARγ-treated FLSs play a pivotal role in OA-related degeneration in ACLT OA rats and IL-1β-treated rat chondrocytes.

**Figure 4.**
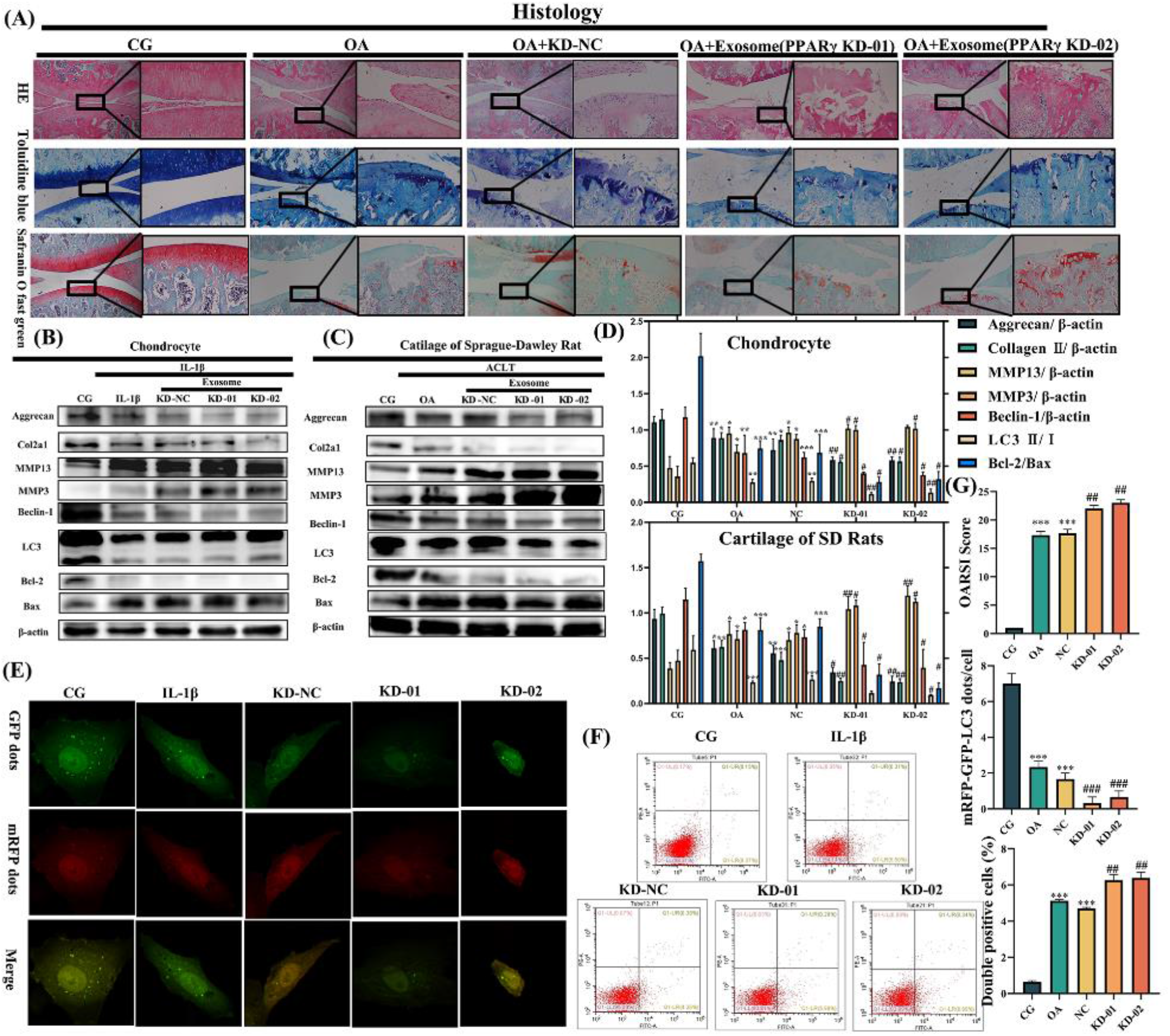
Exosomes derived from KD-PPARγ FLSs exacerbate OA both *in vivo* and *in vitro*. (A) Histological evaluations were performed using hematoxylin & eosin (H&E), toluidine blue, and safranin O-fast green staining. Both KD-01 and KD-02 groups demonstrated aggravated OA symptoms. Relative protein expression was detected in the chondrocytes (B) and in the cartilage of the SD rats (C) that were treated with KD-NC and KD-PPARγ exosome (KD-01, KD-02). (D) Illustrates the ratio of relative protein expression in SD rats and chondrocytes. (E) mRFP-GFP-LC3 adenovirus dual labeling in chondrocytes post OE-PPARγ treatment. mRFP (red) represents autolysosomes (ALs); Merge (yellow) signifies autophagosomes (APs). (F) Flow cytometry analysis shows an increase in chondrocyte apoptosis in the KD-PPARγ group. (G) Presents the OARSI score, the quantification of GFP and mRFP dots per chondrocyte, and the quantification from the flow cytometry analysis. *p < 0.05 vs. CG; **p < 0.01 vs. CG; ***p < 0.001 vs. CG. #p < 0.05 vs. OA; ##p < 0.01 vs. OA; ##p < 0.001 vs. OA. Means ± 95 % CI.

**Figure 5.**
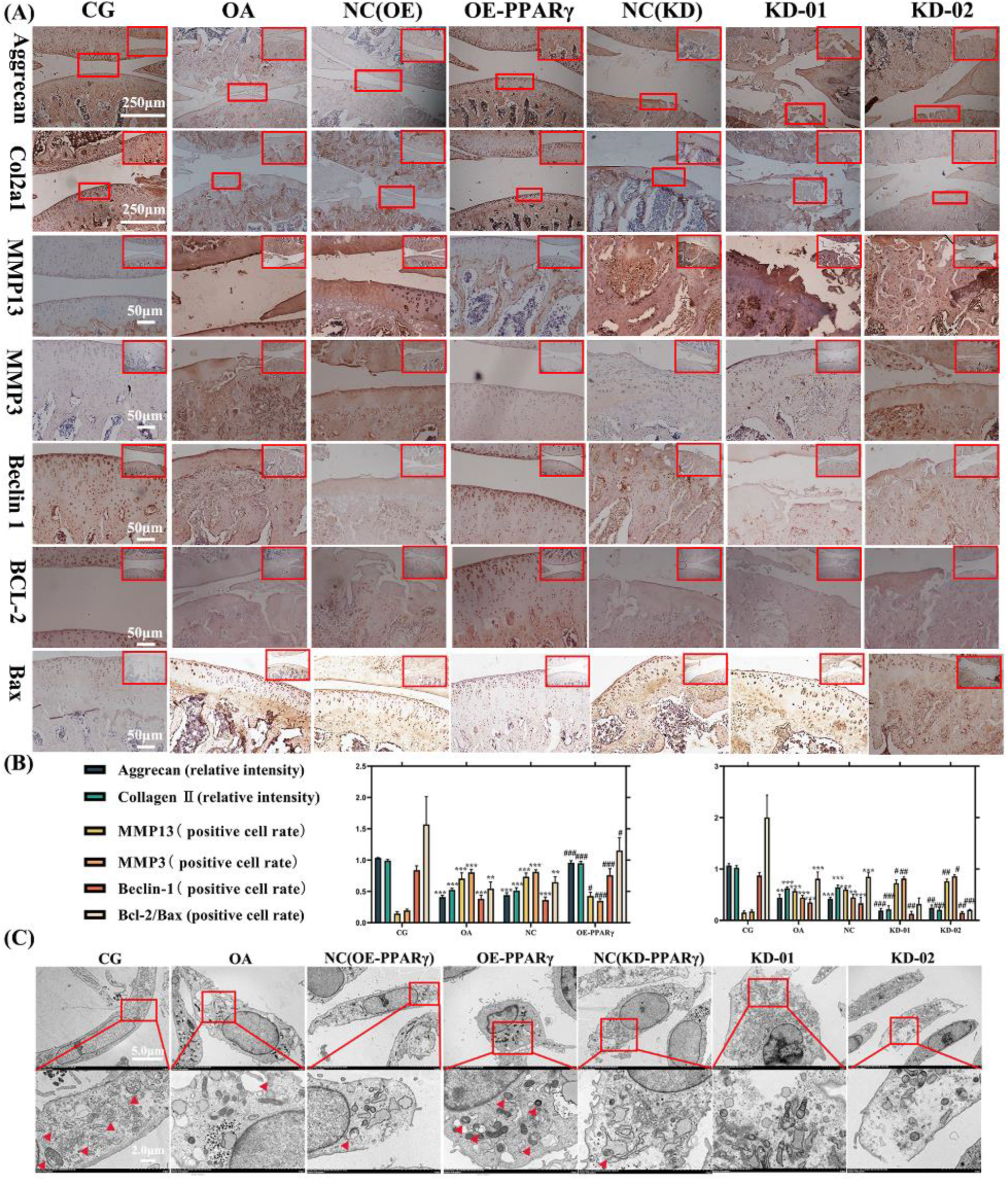
Exosomes derived from OE-PPARγ treated FLSs alleviate OA *in vivo*, while exosomes from KD-PPARγ FLSs exacerbate OA. (A) The cartilage-specific proteins aggrecan and col2α1, which decrease due to OA pathology, are restored by OE-PPARγ exosomes, but are exacerbated by KD-PPARγ exosomes. Inflammatory proteins such as matrix metalloproteinase 13 (MMP13) and 3 (MMP3) are increased in the KD-PPARγ exosome-treated group, but decreased in the OE-PPARγ group compared to the OA group. The treatment with OE-PPARγ exosomes led to an increase in Beclin-1, indicating the activation of autophagy. However, KD-PPARγ (KD-01, KD-02) suppressed this trend. The ratio of BCL-2/Bax was upregulated in OE-PPARγ, but downregulated in KD-PPARγ. (B) The relative intensity of aggrecan, Collagen II, and the positive cell rate of MMP13, MMP3, Beclin-1, BCL-2/Bax were used for immunohistochemistry quantification. Statistical significance is indicated as *p < 0.05 vs. CG; **p < 0.01 vs. CG; ***p < 0.001 vs. CG. #p < 0.05 vs. OA; ##p < 0.01 vs. OA; ###p < 0.001 vs. OA. Means ± 95 % CI. (C) Transmission electron microscopy reveals the ultrastructural features of autophagic vacuoles (APs) in chondrocytes. Red boxes indicate areas of higher magnification, while red arrows point to APs.

### 2.7 Annexin A1 was enriched in OE-PPARγ FLS-derived Exosomes and PPARγ Drove Annexin A1 Expression by PPARγ Coactivator (PGC)-1α

To identify specific proteins within PPARγ-treated FLS-derived exosomes that are involved in treating chondrocyte degeneration, we employed quantitative proteomics to analyze the contents of these exosomes. The Supplementary figure 3 A, B,C illustrate the differentially expressed proteins among the groups, selected according to the criteria of (log2 |fold-change| ≥ 1.2 and p < 0.05). The ANXA1 protein was found to be enriched in the OE-PPARγ group compared with the OE-PPARγ, KO-01, KO-02, and OA groups, as shown in the volcano plot, violin plot, and heatmap analysis.

Upon establishing the potential role of ANXA1 in OE-PPARγ FLS-derived exosomes in treating chondrocyte degeneration, we first utilized Chip-qPCR to investigate the mechanism between PPARγ and ANXA1. However, the Chip results indicated no direct targeting between PPARγ and the ANXA1 promoter (1.22-fold vs the expression of IgG), as depicted in Supplementary figure 2F. Next, we explored the interaction between the PPARγ coactivator PGC-1α and ANXA1 using Co-IP (Figure 6A). The ANXA1 protein was detected in the IP-PGC-1α, and no PGC-1α protein was observed in the band of IgG binding protein. Additionally, IB-PGC-1α was detected in the IP-ANXA1 group. This confirmed the interaction between the PPARγ coactivator PGC-1α and ANXA1, suggesting that ANXA1 may play a crucial role as a mediator in FLS-derived exosomes for the treatment of chondrocyte degeneration.

**Figure 6.**
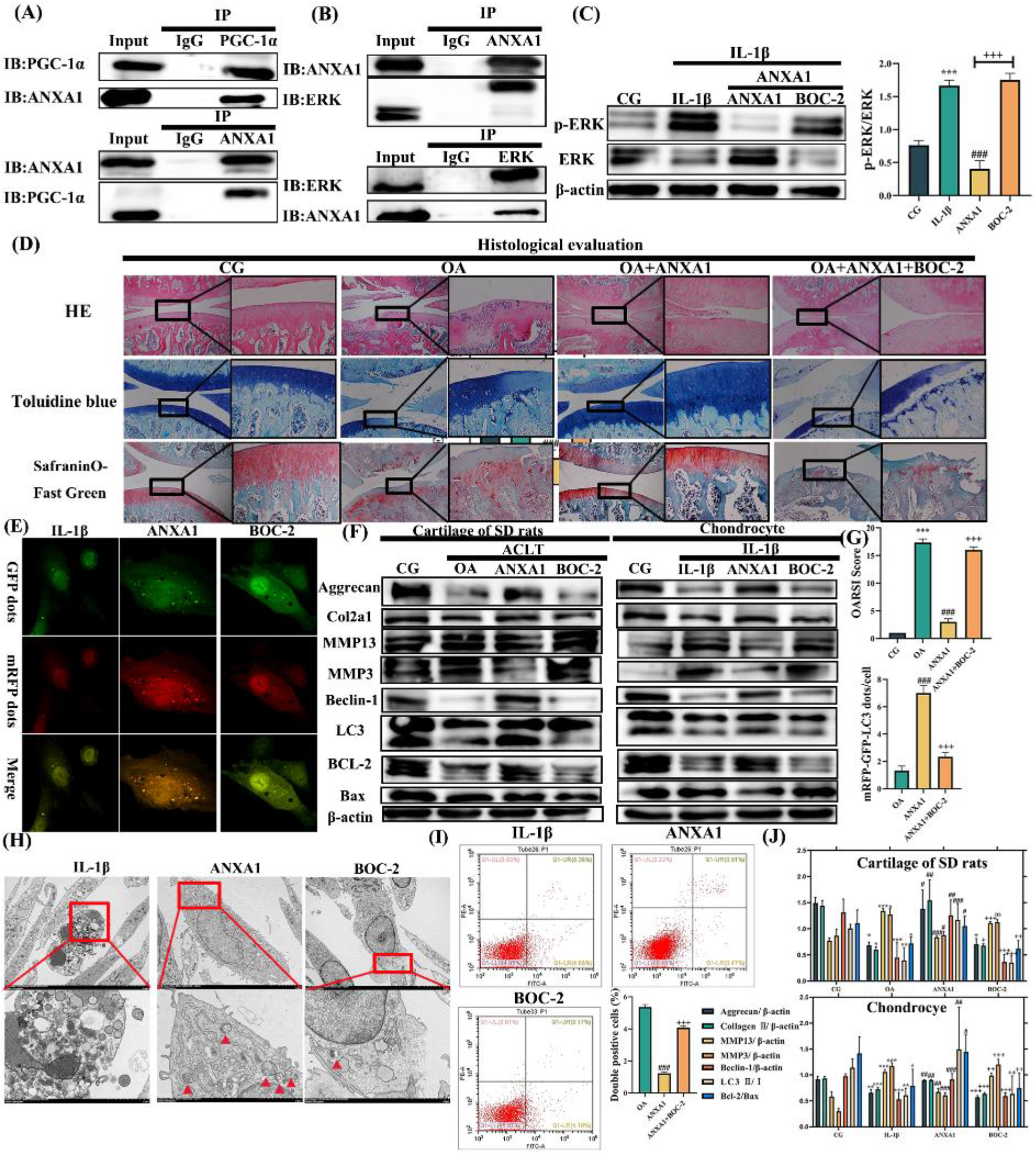
PPARγ stimulates ANXA1 expression via coactivator (PGC)-1α and exosomal ANXA1 alleviates OA-related degeneration *in vivo* and *in vitro* by inhibiting autophagy. Furthermore, ANXA1 can interact with ERK and inhibit its phosphorylation. (A) PGC-1α (upper) and ANXA1 (lower) proteins, immunoprecipitated from FLSs with anti-PGC-1α and anti-ANXA1 antibodies respectively, were analyzed by immunoblotting. (B) ANXA1 (upper) and ERK (lower) proteins, immunoprecipitated from chondrocytes with anti-ANXA1 and anti-ERK antibodies respectively, were also analyzed by immunoblotting. (C) The immunoblotting results (upper) for pERK and ERK, along with their quantification (right), are shown, using β-actin as the endogenous control. (D) Histological analysis showing that the OA+ANXA1 group displays an improvement in OA symptoms, which is reversed by BOC-2. (E) Double labeling of chondrocytes with mRFP-GFP-LC3 adenovirus post-ANXA1 treatment. The ANXA1 group exhibits an increase in autophagosomes (APs), while the BOC-2 group shows a decrease. (F) Relative protein expressions in chondrocytes and cartilage of the SD rats treated with ANXA1 and ANXA1+BOC-2. (H) Transmission electron microscopy illustrates the ultrastructural features of APs in chondrocytes. Red boxes highlight higher magnification regions, with red arrows pointing to APs. (I) Flow cytometry analysis and the ratio of double-positive cells, indicating that ANXA1 reduces chondrocyte apoptosis. (G) The OARSI score and quantification of GFP and mRFP dots per cell in the chondrocytes. (J) The ratio of relative protein expression in SD rats and chondrocytes. *p < 0.05, **p < 0.01, ***p < 0.001 vs. control group (CG); #p < 0.05, ##p < 0.01, ###p < 0.001 vs. OA; +p < 0.05, ++p < 0.01, +++p < 0.001 vs. ANXA1. Means ± 95 % CI. The scale is consistent with previous figures.

### 2.8 Exosomal ANXA1 Ameliorated OA-related degeneration *in vivo* and *in vitro* by Inhibiting Autophagy via Interaction with and Phosphorylation of ERK

Following quantitative proteomics, we utilized GO functional annotation analysis (encompassing Molecular Function, Biological Process, and Cellular Component) and KEGG pathway enrichment analysis to explore potential signaling pathways interacting with ANXA1 released from exosomes. Supplementary figure 4, combined with the bioinformatics analysis of OE-PPARγ versus OA, KO-01, and KO-02 groups, suggests that chondrocyte autophagy, apoptosis, and the ERK cascade are vital in differentially expressed proteins. We then simulated the internalization mechanism of ANXA1 from FLS-derived exosomes by chondrocytes using recombinant rat ANXA1 protein to stimulate chondrocytes. Co-immunoprecipitation (Co-IP) confirmed that ANXA1 is involved in activating the ERK cascade (Figure 6B). We found that ANXA1 intervention could inhibit ERK protein phosphorylation (decreasing the ratio of p-ERK/ERK) compared with the IL-1β group. However, the ANXA1 receptor blocker BOC-2 reversed this inhibitory effect (Figure 6C).

Next, we employed western blotting (Figure 6F), mRFP-GFP-LC3 adenovirus double labeling (Figure 6E), and transmission electron microscopy (Figure 6H) to determine if ANXA1 regulates autophagy by phosphorylating ERK. Our results showed that expression levels of aggrecan and Col2α1 proteins, downregulated by IL-1β and the OA model, were reversed after ANXA1 treatment. MMP-13 and MMP-3 were decreased in the ANXA1 group. Treatment with ANXA1 increased Beclin-1 and LC3Ⅱ/Ⅰ levels, indicating autophagy activation. The ratio of Bcl-2 and Bax was upregulated in ANXA1-treated samples, suggesting that ANXA1 could reduce chondrocyte apoptosis. Furthermore, mRFP-GFP-LC3 adenovirus double labeling and TEM showed autophagosomes (APs) of chondrocytes were upregulated by ANXA1. Flow cytometry analysis results (Figure 6I) also showed that apoptosis decreased after intervention with ANXA1. However, these effects were reversed by the ANXA1 receptor blocker BOC-2. *In vivo*, we used histological evaluation (Figure 6D), immunohistochemistry (Figure 7A), and western blotting of the cartilage in SD rats to evaluate these parameters. Collectively, these results suggest that ANXA1 ameliorates OA-related degeneration by interacting with and phosphorylating ERK, thereby activating autophagy.

**Figure 7.**
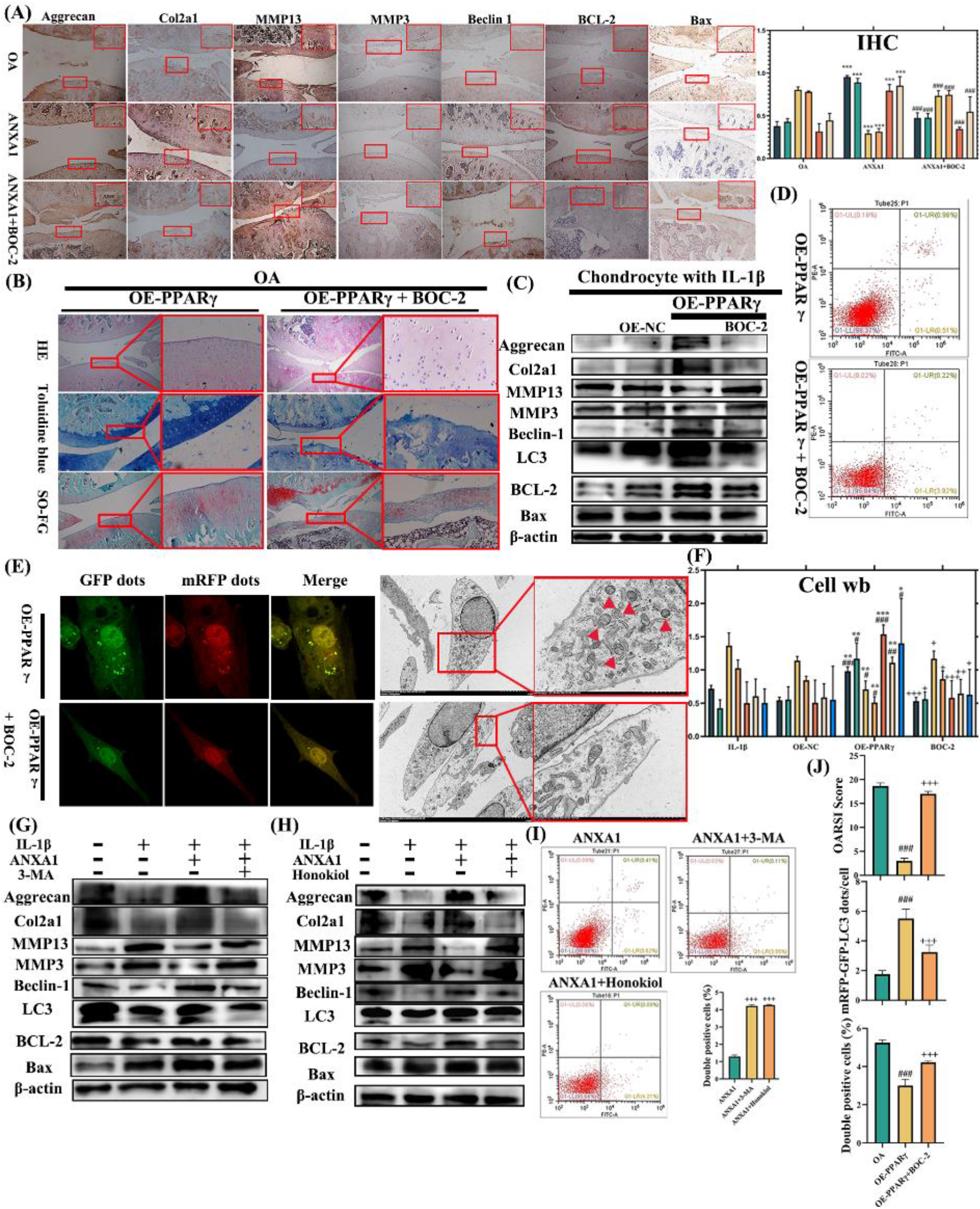
Rescue experiment of the PPARγ – FLSs exosomal ANXA1 – ERK axis in OA. (A) Depicts the results of Immunohistochemistry (IHC) for ANXA1 and BOC-2 treatment *in vivo*. (B) Histological analysis illustrating that the therapeutic effect of overexpressed (OE) PPARγ exosomes can be blocked by the ANXA1 receptor blocker BOC-2. (C) Demonstrates the relative protein expression levels in chondrocytes treated with OE-PPARγ exosomes and BOC-2. (D) Flow cytometry analysis showing that BOC-2 reverses the therapeutic effect of OE-PPARγ exosomes. (E) Double labeling with mRFP-GFP-LC3 adenovirus in chondrocytes, revealing that BOC-2 treatment reduces autophagosomes (APs). (E) Transmission electron microscopy images highlighting the ultrastructural features of APs in chondrocytes. The upper group is treated with OE-PPARγ exosomes, and the lower group with BOC-2. (G) Western blot analysis of chondrocytes treated with the autophagy inhibitor 3-MA, indicating that 3-MA diminishes the therapeutic effect of ANXA1. (H) Western blot analysis of chondrocytes treated with ANXA1 and the ERK activator Honokiol, demonstrating that the Honokiol group shows a decreased therapeutic effect of ANXA1. (J) The OARSI score, quantification of GFP and mRFP dots per cell in chondrocytes, and quantification of flow cytometry analysis.

**Figure 8.**
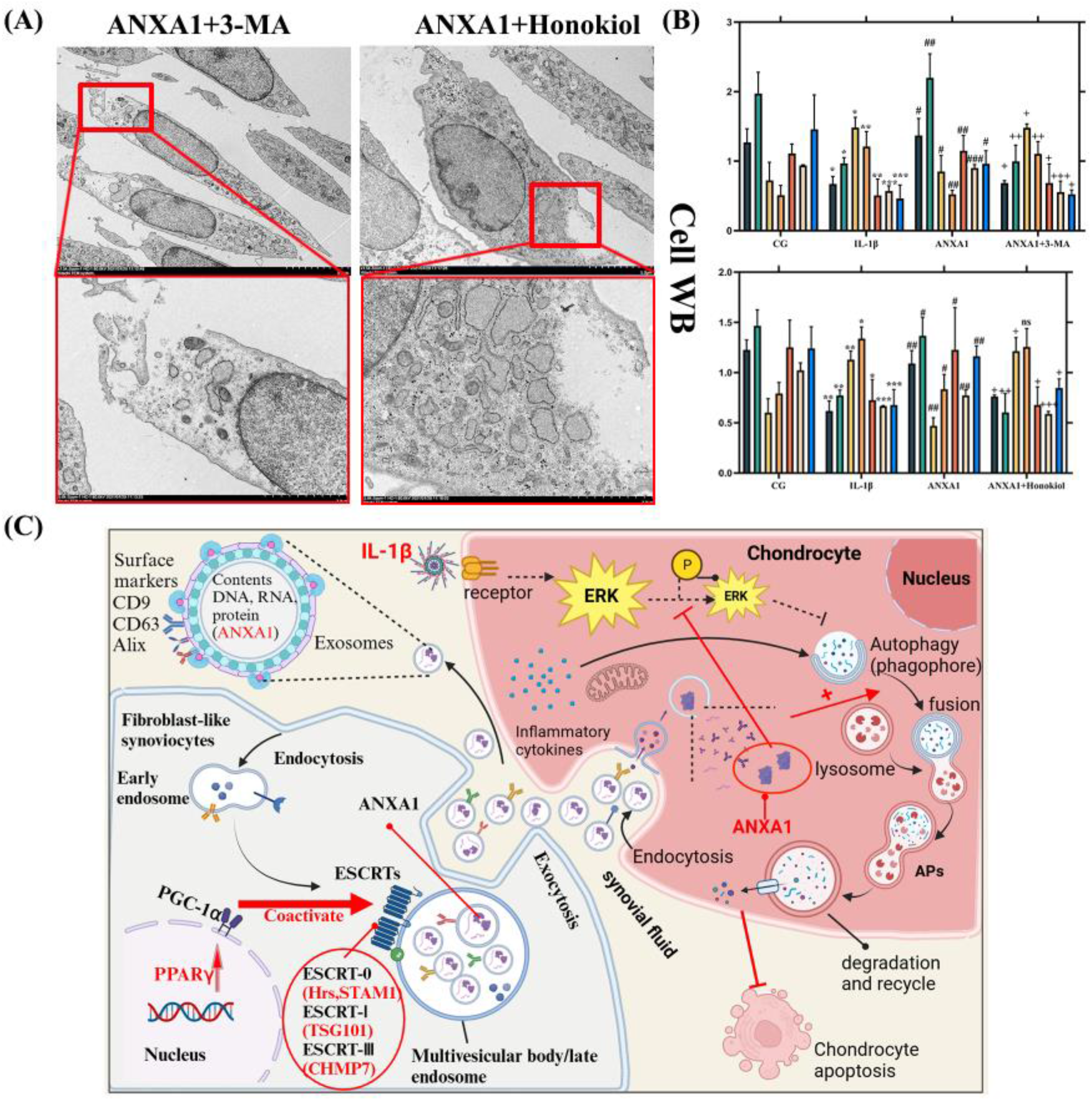
PPARγ controls ESCRT-dependent fibroblast-like synoviocyte exosome biogenesis and Alleviates Chondrocyte Osteoarthritis Mediated by Exosomal ANXA1.

### 2.9 PPARγ-exomal ANXA1-ERK Axis Participated in the Progression of OA Cartilage Degeneration by Regulating Autophagy

To further explore the exercise-induced PPARγ-ANXA1 (derived from FLSs exosomes)-ERK axis, we utilized a rescue experiment to validate the significance of this axis in OA’s exercise therapy. Initially, chondrocytes were treated with overexpressed (OE) PPARγ FLS-derived exosomes and the ANXA1 receptor blocker BOC-2. Following histological evaluation (Figure 7B), western blot analysis (Figure 7C, F), mRFP-GFP-LC3 adenovirus double labeling (Figure 7E, J), TEM (Figure 7E), and flow cytometry analysis (Figure 7D, K), we found that BOC-2 intervention inhibits the therapeutic effect of exosomes, thereby decreasing autophagy and increasing chondrocyte apoptosis. This suggests that the upregulation of PPARγ in FLSs influences chondrocytes via ANXA1 in exosomes.

To ascertain the role of autophagy in the ANXA1 treatment of OA, we compared the ANXA1+3-MA group (with autophagy inhibitor) to the ANXA1 group (Figure 7G, I). We noted that the apoptosis, MMP13, and MMP3 levels reduced by ANXA1, as well as the upregulation of COL2α1 and aggrecan, were partially reversed after treatment with the autophagy inhibitor 3-methyladenine (3-MA). We hypothesized that ANXA1 intensifies degeneration by reducing autophagy in chondrocytes. Moreover, the ANXA1-induced upregulation of COL2α1, aggrecan, autophagy markers Beclin-1 and LC3 Ⅱ/Ⅰ, BCL-2/Bax, and the decrease of MMP13, MMP3 levels were reversed following treatment with the ERK activator Honokiol (Figure 7H) (Supplementary figure 1F).

Collectively, these findings suggest that PPARγ in FLSs stimulates the release of ANXA1 in exosomes through ESCRT-dependent pathway, which then interacts with and inhibits ERK phosphorylation. This interaction results in the activation of autophagy, thereby mitigating OA degeneration.

## 3. Discussion

This study highlighted exosomes derived from FLSs as a novel communication factor between FLSs and chondrocytes in exercise therapy for OA. We firstly reported that PPARγ regulated exosome biogenesis and cargo exercise related proteins through ESCRT-dependent pathway. And our study proposes a therapeutic mechanism whereby the exercise-associated gene PPARγ, induces the generation of FLS-derived exosomes and regulates exosomal protein content (ANXA1) in OA through cell communication between FLSs and chondrocytes. We also observed that OE-PPARγ in FLS-derived exosomes could alleviate chondrocyte apoptosis, whereas KD-PPARγ in FLS-derived exosomes could exacerbate OA. Furthermore, our findings showed enrichment of ANXA1 in the OE-PPARγ FLS-derived exosomes and confirmed that the therapeutic agent in the exosome is indeed ANXA1. Finally, we concluded that the exercise-induced PPARγ/ESCRT – FLSs exosomal ANXA1 – ERK axis mitigates OA through the activation of autophagy.

We established OA models *in vivo* and *in vitro* using the ACLT surgical technique [39] and IL-1β [9, 31]. We introduced a moderate treadmill exercise regime into the OA model [9, 10] and utilized single-cell transcriptome sequencing to analyze the model’s synovium, cartilage, and subchondral bone, which correspond to the most significant pathological features of OA: synovitis, cartilage degeneration [40].

In articular joints, the microenvironment of cartilage is made up of synovial fluid (SF). FLSs in the synovium are responsible for producing the viscous SF that fills the joint cavity [41]. SF not only lubricates the articular cartilage and serves as a medium for nutrient and waste transportation, but its role in mediating intercellular communication through soluble factors has gained increasing attention in recent years [42]. Under pathological or physiological conditions, changes in the molecular and cellular components of SF can directly impact articular cartilage. This unique characteristic positions FLSs and their secretions as an intriguing source for OA therapy. Therefore, in isolated FLSs, we found 247 genes that were upregulated in the exercise plus OA group compared to the sedentary OA group (Supplementary figure 1A-E). Previous studies [43] have primarily focused on the role of PPARγ in cartilage, but have overlooked the influence of PPARγ, widely believed to be upregulated by exercise, as an intercellular communicator within the joint. We hypothesized that cytokines secreted by PPARγ-treated synovium might play a role in cartilage treatment. The co-culture of PPARγ-treated FLSs and chondrocytes supported this hypothesis. Subsequently, we investigated the specific mechanism of communication between FLSs and chondrocytes.

The synovium, a secretory tissue, is abundant in soluble inflammatory mediators and extracellular vesicles, including exosomes. Exosomes participate in various physiological processes such as apoptosis, angiogenesis, inflammation, and coagulation. They also transport cargo like proteins, lipids, and RNA, thereby regulating cellular communication and epigenetic modifications [44, 45]. However, the molecular mechanisms of exosome biogenesis has been less intensively investigated. Exosome biogenesis consists of four steps: ILVs formation, cargo sorting to MVBs, MVB formation, transport of MVBs [18, 19]. ESCRT is well studied for its function in membrane budding and ILVs formation. ESCRT consists of ESCRT-0, -I, -II, -III subcomplexes [46]. The ESCRT-0 complex is responsible for the initial recognition of ubiquitinated protein cargo and initiation of the ESCRT pathway by the recruitment of ESCRT-I and –II. ESCRT-0 contains two subunits, hepatocyte growth factor-regulated tyrosine kinase substrate (HRS), and signal-transducing adaptor molecule 1 (STAM 1) [24]. Then, ESCRT-I (TSG101) and –II cooperate to form a saddle-shaped protein complex that is important for the ESCRT-III assembly [21]. ESCRT-III (CHMP7) is involved in membrane deformation, facilitate inward budding and vesicle abscission to form ILVs [46]. Therefore, the ESCRT machinery plays a key role in exosome biogenesis. We found PPARγ and PGC-1α could interacted with ESCRT-0 (Hrs and STAM1), ESCRT-I (TSG101) and ESCRT-III (CHMP7) subunits and regulated their expression. Additionally, we also observed that PPARγ colocalized with the endosomal vesicle structures which indicate exosome endocytosis pathway [47]. Taken together, we demonstrate exercise-PPARγ controls ESCRT-dependent exosome biogenesis which carry unique cargo.

We isolated and characterized exosomes derived from OE-PPARγ and KD-PPARγ FLSs. Although the exosomes we isolated were slightly larger in diameter than those reported in previous studies, their morphology was comparable [16, 48]. Prior research has established that the expression of a single exosomal cargo protein can increase exosome density and significantly alter their size and shape. As such, exosome size, shape, and density are highly variable and do not necessarily define exosomes [16, 42]. When incubated with differentially treated FLS-derived exosomes, the OE-PPARγ group showed increased autophagy, decreased apoptosis, and thus ameliorated chondrocyte degeneration. Conversely, the KD-PPARγ exosomes reversed these therapeutic effects. *In vivo*, OE-PPARγ exosome treatment reduced osteophyte formation and cartilage destruction. The beneficial role of FLS-derived exosomes is further demonstrated by their capacity to initiate the cartilage repair response, facilitate matrix deposition, and promote tissue formation. Thus, we confirmed that exosomes derived from PPARγ-treated FLSs participate in the pathological processes affecting cartilage and chondrocytes in OA. Given that exosomes have been found to carry numerous types of proteins involved in disease development and prognosis [49], and their composition reflects the cellular source from which they originate, better characterization of exosomal cargoes may facilitate the identification of new indicators and inform targeted therapy for OA.

Data-independent acquisition (DIA) represents a recently developed mass spectrometry data collection method [50]. We utilized DIA quantitative proteomics technique to examine the protein profiles of PPARγ-treated FLSs-derived exosomes. The findings demonstrated 27 upregulated and 55 downregulated proteins in the OE vs KD-01 comparison; 70 upregulated and 147 downregulated proteins in the OE vs KD-02 comparison; and 42 upregulated and 58 downregulated proteins in the OE vs OA comparison. Across these comparisons, ANXA1 was found to be enriched in the OE-PPARγ group compared to the KD-PPARγ and OA groups. ANXA1 has been identified as a constituent of exosomes or extracellular vesicles [51, 52]. ANXA1 significantly influences many physiological processes such as cell growth, differentiation, membrane fusion, endocytosis, and exocytosis [53, 54]. Anti-inflammatory effects and enhanced autophagic flux have been reported, with ANXA1 needing to be externalized to the cell membrane or secreted into the extracellular fluids to exert its anti-inflammatory properties [55, 56]. Considering that the exogenous intervention of ANXA1 can also be mediated by the cell surface receptor for annexin A1, N-formyl peptide receptor 2 (FPR2, also known as lipoxin A4 receptor or ALX) [57], we stimulated OA chondrocytes with recombinant ANXA1 to mimic exosomal ANXA1 internalization by chondrocytes. Our results indicated that ANXA1 intervention could increase autophagy flux and decrease apoptosis both *in vivo* and *in vitro*. The ANXA1 receptor antagonist, BOC-2, could reverse this effect. Nonetheless, ANXA1 has been reported to exhibit both anti- and pro-inflammatory effects [58, 59], acting as a double-edged sword. Its effects are largely mediated through formyl peptide receptors (FPRs), which bind a wide range of ligands [60]. The multifaceted nature of ANXA1 ligands, the multiple signaling pathways, and the molecule’s localization perhaps contribute to this disparity [61]. To validate the anti-inflammatory effect of ANXA1 on OA chondrocytes, the upstream and downstream ligands in chondrocytes that interact with ANXA1 need to be defined.

In our study, we established that the promoter of ANAX1 is not directly regulated by PPARγ, as evidenced by Chip-qPCR experiments (see Supplementary figure 2F). However, our Co-IP results demonstrated that the coactivator PPARγ coactivator-1α (PGC-1α) promotes ANXA1 expression (Figure 6A). Hence, PPARγ enhances the expression of ANXA1 in exosomes via its coactivator PGC-1α, aligning with previous research [30]. To clarify the precise molecular mechanism governing ANXA1’s regulation of cartilage degeneration, we identified apoptosis, autophagy, and the ERK cascade as potential cellular processes and components of environmental information processing, using KEGG and GO analysis. Co-IP results indicated that ANXA1 can interact with the ERK protein (Figure 6B, C), and ANXA1 intervention can inhibit the phosphorylation of the ERK cascade. Notably, BOC-2 also reversed this trend. The finding that external treatment with ANXA1 can suppress ERK phosphorylation is intriguing, considering that the activation of this pathway can inhibit autophagy and increase apoptosis [31–33]. Consistent with this, autophagy stimulated by ANXA1 was reversed by the ERK activator Honokiol and the autophagy inhibitor 3-MA. This suggests that ANXA1 inhibits autophagy by interacting with and phosphorylating ERK.

Our study has several limitations. First, while we have identified the potential mechanism through which exercise-induced PPARγ-FLSs exosomal ANXA1-ERK axis regulates autophagy to mitigate chondrocyte degeneration, there are numerous potential genes and binding partners in this axis that have not been explored. Further study is required to fully understand this relationship. Second, we manipulated ANXA1 directly using recombinant ANXA1 protein, rather than constructing exosomes coated with ANXA1. Third, the intra-articular injection of the KD-PPARγ adenovirus, used in our study, doesn’t only affect synovial tissue but also impacts cartilage tissue. Therefore, we did not conduct a rescue experiment of exercise combined with synovial KD-PPARγ.

In conclusion, our study revealed that moderate exercise enhances the expression of PPARγ in fibroblast-like synoviocytes (FLSs). Moreover, exercise-induced PPARγ and PGC-1α in FLSs can augment the content of the ANXA1 protein in exosomes derived from FLSs through ESCRT-dependent exosome biogenesis pathway. This exosomal ANXA1 subsequently inhibits ERK phosphorylation, thereby activating chondrocyte autophagy. Importantly, we suggest that the exercise-induced PPARγ/ESCRT – FLSs exosomal ANXA1 – ERK axis contributes to alleviating OA cartilage degeneration by inhibiting apoptosis. By investigating the interaction between FLSs and chondrocytes, profiling exosomal cargo, and understanding the mechanisms affecting cartilage, our study could unveil novel regulatory mechanisms within the joint. This insight may lead to more targeted treatment strategies for OA.

## 4. Materials and Methods

### 4.1 Experimental Animals and Osteoarthritis Model

Male Sprague–Dawley (SD) rats (240 ± 5 g, 4-week-old, specific-pathogen-free, eight rats per group, and six rats per cage) were procured from HFK Bioscience Co. Ltd. (Beijing, China). The study was approved by the Ethics Committee of China Medical University (no. 2017PS237K), adhering to the regulations stipulated in the World Medical Association Helsinki Declaration. The upkeep and care of the experimental rats complied with the committee’s guidelines, as described in previous studies [9, 10].

The OA model was established using Anterior Cruciate Ligament Transection (ACLT) (refer to Supplementary figure 2A) [39]. The surgical procedure involved a 3 mm longitudinal incision from the distal patella to the proximal tibial plateau. The medial capsula articularis, adjacent to the patellar tendon, was incised with a scalpel, and the capsula articularis was further incised with microiris scissors. The fat pad was carefully removed from the intercondylar area, revealing the anterior cruciate ligament (ACL). The ACL, which extends from the distal end of the femur and attaches to the anteromedial portion of the tibia, was exposed by dislocating the patella medially and keeping the cartilage moist with saline. The ACL was then transected with a micro-surgical knife under direct visualization and confirmed by an anterior drawer test. The incision was subsequently sutured layer by layer. The model was established over a period of 4 weeks, following previous studies [39]. The 36 rats were segregated into three groups: the control group (CG), the osteoarthritis group (OA), and the osteoarthritis plus moderate exercise group (EXE+OA), with 12 rats in each group. Post-modelling, the EXE+OA group began treadmill training on the ZH-PT animal treadmill exercise platform (Zhongshidichuang Science & Technology Development Co. Ltd., Beijing, China) using appropriate light, acoustic, and electrical stimuli. The moderate treadmill exercise protocol (19.2 m/min) was based on our prior studies [9, 10, 12, 62]. As our previous research demonstrated that moderate-intensity exercise has the optimal therapeutic effect on osteoarthritis [9–13, 63], we excluded low and high-intensity exercise from this study. A total of 36 SD rats were used for single-cell transcriptome sequencing (n=12).

### 4.2 Tissue Sampling and Collection

After the final treadmill exercise session, rats were euthanized via a pentobarbital overdose, as per our established protocol [9, 10]. Tissue samples, including synovium, cartilage, and subchondral bone, were subsequently collected for single-cell transcriptome sequencing.

For histological and immunohistochemical evaluations, we employed an additional 54 SD rats. Their entire knee joints were fixed using a 4% paraformaldehyde solution (Sigma-Aldrich, St. Louis, MO, United States) for a duration of 7 days. Subsequently, the samples underwent decalcification in a 20% EDTA solution (Sigma-Aldrich, St. Louis, MO, USA) for 7 weeks. Following decalcification, the samples were dehydrated using a graded series of ethanol and xylene (Sigma-Aldrich) and embedded in paraffin (Sigma-Aldrich) [9].

### 4.3 Single-cell Transcriptome Sequencing and Bioinformatics Analysis

Synovium, cartilage, and subchondral bone were collected from the CG, OA, and EXE+OA groups. Sequencing libraries were generated following a comprehensive transcriptome sequencing and single-cell gene-expression profiling protocol as previously reported [64, 65]. In summary, tissues were cleansed in ice-cold RPMI1640 and dissociated using the Multi-tissue dissociation kit 2 (Catalog No.130-110-203, Miltenyi Biotec, USA) following the manufacturer instructions. Depending on the viscosity of the homogenate, DNase treatment was optionally applied. Cell count and viability were evaluated using a fluorescence Cell Analyzer (Countstar® Rigel S2, Alit Biotech, Shanghai, China) with AO/PI reagent after erythrocyte removal (Miltenyi 130-094-183). Debris and dead cells were then eliminated (Miltenyi 130-109-398/130-090-101). Finally, fresh cells were washed twice in RPMI1640 and resuspended at 1×10^6^ cells/mL in 1× phosphate-buffered saline (PBS) with 0.04% bovine serum albumin.

Single-cell RNA-Seq libraries were generated using the SeekOne® Digital Droplet Single Cell 3’ library preparation kit (SeekGene, Beijing, China). Briefly, an appropriate number of cells were mixed with reverse transcription reagent and added to the sample well in SeekOne® DD Chip S3. Barcoded Hydrogel Beads (BHBs) and partitioning oil were then dispensed into the corresponding wells in chip S3. After emulsion droplet generation, reverse transcription was performed at 42℃ for 90 min and inactivated at 80℃ for 15 min. Next, cDNA was extracted from the ruptured droplets and amplified in a PCR reaction. The amplified cDNA product was cleaned, fragmented, end-repaired, A-tailed, and ligated to a sequencing adaptor. The indexed PCR was performed to amplify the DNA representing the 3ʹpolyA part of expressing genes, which also contained the Cell Barcode and Unique Molecular Index. The indexed sequencing libraries were cleansed with SPRI beads, quantified by quantitative PCR (KAPABiosystems KK4824) and sequenced on the Illumina NovaSeq 6000 with PE150 read length or DNBSEQ-T7 platform with PE100 read length.

To identify exercise-related differentially expressed genes in FLSs, we compared the expression profiles of the FLSs of the CG, OA, and EXE+OA groups. Volcano plots of differentially expressed genes were generated using the Limma/R package (version 3.5.1). The primary parameters were set as log fold change, with a log fold change >2 indicating differentially expressed genes. These differentially expressed genes were then submitted to the DAVID v6.8 tool (https://david.ncifcrf.gov/) for annotation, visualization, and integrated discovery [9]. Gene ontology (GO) functional annotation (covering Molecular function, Biological process, and Cellular component) and Kyoto Encyclopedia of Genes and Genomes (KEGG) pathway enrichment analyses were used to identify potential pathways.

### 4.4 Histological Examination of Articular Cartilage in Human and Animal Models

The study protocol and experimental procedures involving joint specimens obtained from human arthroscope or total knee arthroplasty were approved by the Ethics Committee of Shengjing Hospital of China Medical University (Approval No: 2019PS629K), in accordance with the principles of the World Medical Association’s Helsinki Declaration. The synovium and clinicopathological features of patients with different Kellgren-Lawrence (K-L) grades were collected, based on data from 114 patients treated by arthroscope or arthroplasty in our hospital. Donors participated in this study, and informed consent was obtained from all participants. The inclusion criteria and exclusion criteria are described as our previous study [66]. As detailed in our earlier research [9, 31], we categorized the synovium and the clinicopathological characteristics of patients with knee osteoarthritis into with different K-L grades. This categorization was guided by the Osteoarthritis Research Society International (OARSI) scale, which uses a point system ranging from 0 to 4 [67].

Following this, we embedded joints from SD rats in paraffin. Sagittal sections of 4.5-μm thickness were cut from the tibiofemoral joints and stained with hematoxylin and eosin (H&E), toluidine blue, and safranin O-fast green (SO-FG) for histological evaluation as described in [9, 11]. The degree of articular cartilage damage in the tibiofemoral joint was assessed using the OARSI score [67]. This scoring was carried out by two experienced observers who were blind to the study group allocations.

### 4.5 Primary Chondrocyte and FLS Isolation, Culture, and Treatment

Synovial tissue and articular cartilage were harvested from SD rats. The synovium was digested with 1 mg/mL collagenase (SigmaAldrich) for 2–3 h at 37°C [12]. Articular cartilage samples were digested in pronase (2 mg/mL, Roche, Basel, Switzerland) and collagenase D (1 mg/mL; Roche, Basel, Switzerland) [9]. The extracted cells were cultured as previously detailed [9, 12]. To maintain the phenotypic integrity of the primary FLSs and chondrocytes, and to ensure the accuracy of subsequent experiments, only 2–3 passages were utilized. An *in vitro* OA model was induced by treating rat primary chondrocytes with recombinant rat Interleukin-1β (IL-1β) (10 ng/mL) (MedChemExpress, Shanghai, China) for 24 h, as per previously described protocols [9, 31].

The role of PPARγ in OA treatment was assessed by transfecting primary FLSs with a recombinant adenovirus carrying either PPARγ knockdown or overexpression sequences (multiplicity of infection [MOI] = 50, polybrene = 1 μg/mL; Hanbio, China). Post transfection, the FLSs were subjected to IL-1β stimulation (10 ng/mL) for 24 h. PPARγ knockdown conditions were grouped as follows: IL-1β + sh-NC (KD-NC), IL-1β + HBAD-Adeasy-r- PPARγ shRNA1-EGFP (KD-01), and IL-1β + HBAD-Adeasy-r- PPARγ shRNA2-EGFP (KD-02). PPARγ overexpression conditions were divided into IL-1β + Ad-NC (OE-NC) and IL-1β + HBAD-Adeasy-r- PPARγ-3xflag-EGFP (OE-PPARγ). In addition, primary chondrocytes *in vitro* were treated with exosomes derived from FLSs, and the experimental conditions were organized into the following groups: CG, IL-1β, OE-NC, OE-PPARγ, KD-NC, KD-01, KD-02.

To examine the role of ANXA1 in regulating autophagy via ERK phosphorylation *in vitro*, chondrocytes were treated with 130 ng/mL rat ANXA1 protein (Recombinant His-S) (G15574, LifeSpan BioSciences, Seattle, WA, USA) and simultaneously stimulated with IL-1β (10 ng/mL) and the ANXA1 receptor antagonist BOC-2 [68, 69] for 24 h. The concentration of administered ANXA1 protein was determined based on quantitative proteomics (Section 2.9) of exosomal ANXA1. These experimental conditions were divided into IL-1β + ANXA1 (ANXA1), and IL-1β + ANXA1 + BOC-2 (ANXA1+BOC-2) groups. Subsequently, the ERK activator Honokiol (5 μM; MedChemExpress) and the autophagy inhibitor 3-methyladenine (3-MA; 5 mM; MedChemExpress) were applied to validate their respective effects.

### 4.6 Co-culture of PPARγ-treated FLSs with Primary Chondrocytes

To mimic the physiological interaction between the synovium and cartilage, we established a non-contact co-culture system using a Transwell apparatus (Model No: 3450, pore size: 0.4 μm; Corning Costar Corp., NY, USA), as described in a previous study [70]. Chondrocytes (5×10^4^ cells/well) were placed in the bottom compartment of the Transwell system, while PPARγ-treated FLSs (5×10^4^ cells/well) were positioned in the top compartment. Cells were cultured using 0.4 µm inserts, enabling communication between FLSs and chondrocytes via a polyester (PET) membrane for a period of 48 h. After this incubation period, the chondrocytes were harvested for subsequent investigations [70].

### 4.7 Isolation, Characterization, and Tracking of FLS-derived Exosomes

Primary FLSs were transfected with recombinant adenovirus for either PPARγ knockdown or overexpression. Exosomes were then isolated from the culture medium through ultracentrifugation [71, 72]. Initially, the culture medium was centrifuged at 300 ×*g* for 5 min to remove dead cells. The remaining supernatant was further centrifuged at 2,000 ×*g* for 15 min, followed by centrifugation at 13,000 ×*g* for 35 min to eliminate macromolecules. After filtration through a 0.22 μm filter membrane, the supernatant was transferred to an ultrafiltration tube for centrifugation at 100,000 ×*g* for 70 min. The resulting supernatants, referred to as exosome-depleted conditioned media (CM-exo), were collected, and the purified exosomes were resuspended in DMEM for direct use in subsequent experiments.

Transmission electron microscopy (TEM) was used to clearly visualize the typical structure of exosomes. Exosome samples were removed from the -80℃ refrigerator, placed in an ice box, and gently centrifuged after thawing. Fifteen microliters of exosome samples were pipetted onto a copper grid for one minute. After blotting with filter paper, the samples on the copper grid were stained with 15μL of 2% uranyl acetate for one minute and air-dried under a lamp for 10 min. The samples were subsequently observed and imaged using a Tecnai G2 Spirit TEM (FEI, USA).

Nanoparticle Tracking Analysis (NTA), a recognized technique for characterizing exosomes, was used to track and analyze the Brownian motion of each particle. This allowed the calculation of the hydrodynamic diameter and concentration of nanoparticles, according to the Stokes–Einstein equation [73]. The number and size distribution of exosomes were analyzed using the Nanosight NS300 system (Nanosight Technology, Malvern, UK), following the manufacturer’s instructions. Western blot analysis was performed to detect the presence of exosome markers (CD9, CD63, ALIX). Calnexin was used for exosome negative control

To explore whether FLS-derived exosomes could be internalized by chondrocytes, the exosomes were labeled with PKH-67 (green, HY-D1421, MedChemExpress) and PKH-26(red, HY-D1451, MedChemExpress). Primary chondrocytes were stained with 4,6-diamidino-2-phenylindole (DAPI) for 5 min and visualized under a confocal microscope (Olympus).

### 4.8 Treatment of OA model with FLS-derived Exosomes and ANXA1 Protein

Exosomes were isolated from seven different groups: CG, IL-1β, OE-NC, OE-PPARγ, KD-NC, KD-01, and KD-02. To explore the role of PPARγ, exosomes derived from these groups were administered into the intra-articular cavity of SD rats at a dose of 4×10^6^ particles per rat, thrice weekly before ACLT [74, 75]. Consequently, 42 SD rats, treated with PPARγ-modified FLS-derived exosomes, were divided into seven groups (n = 6 per group): CG, OA, OA + OE (NC), OA+ HBAD-Adeasy-r-PPARγ-3xflag-EGFP (OE-PPARγ), OA+sh-NC (KD (NC)), OA + sh-PPARγ-01 (KD-01), and OA + sh-PPARγ-02 (KD-02). To discern the role of ANXA1, recombinant His-S ANXA1 protein (G15574, LifeSpan BioSciences, Seattle, WA, USA) was injected into the intra-articular cavity of SD rats four times weekly (1.3μg/50μL) before implementing ACLT in the OA model. Rats that received ANXA1 and the ANXA1 receptor antagonist BOC-2 [68, 69] were divided into two groups (n = 6 per group): ANXA1 and ANXA1+BOC-2 group. Altogether, 54 SD rats were used for evaluating PPARγ and ANXA1 (n=6).

### 4.9 Quantitative Proteomics of FLS-derived Exosomes via Data-independent Acquisition and Related Bioinformatics Analysis

Following previous studies [50, 76], all exosomes were enzymatically digested with trypsin. Subsequently, iodoacetamide was added, followed by six volumes of pre-cooled acetone to precipitate the protein. The precipitated protein was then redissolved with an enzymatic diluent [protein: enzyme = 50:1 (m/m)] and lyophilized.

To examine the protein profiles of the exosomes, we employed the Tandem Mass Tags (TMT)-labeled quantitative proteomics technique [76]. All analyses were conducted using a Q-Exactive HFX mass spectrometer (Thermo, USA) equipped with a Nanospray Flex source (Thermo, USA). Chromatographic separation was carried out on the EASY-nLC 1000 HPLC System (Thermo, USA). Following each full MS scan, 20 MS2 scans were collected as per the inclusion list. The detection mode was positive ion, with a primary MS scanning range of 350-1650 m/z, an MS resolution of 60,000 (m/z 200), an AGC target of 3e6, and a maximum IT of 50ms. MS2 data was acquired using the DIA mode, setting 30 DIA acquisition windows, mass spectral resolution: 30,000 (m/z 200), AGC target:3e6, maximum IT: auto, MS2 activation type: HCD, normalized collision energy: 30, and spectral data type: profile.

All of the Q Exactive raw data was searched using DIA-NN (v1.8.1) and the Ensembl Rattus 45936 20220121.fasta database. A global false discovery rate (FDR) was set to 0.01, and protein groups were considered for quantification if they had at least 2 peptides. Finally, the differences between each group were analyzed using bioinformatics analysis (as mentioned above).

### 4.10 Western Blot Analysis

Both cartilage and primary chondrocytes were lysed using RIPA (9806S, Cell Signaling Technology), supplemented with 1 mM PMSF (ST506; Beyotime Biotech, Shanghai, China) and 1 mM phosphatase inhibitors (P1081; Beyotime Biotech, Shanghai, China). Equivalent amounts of proteins (40 μg) were separated by polyacrylamide gel electrophoresis (8–15% SDS-PAGE) and subsequently transferred onto polyvinylidene difluoride (PVDF) membranes. The subsequent steps adhered to our previously established protocol [9, 31, 66]. The antibodies used included anti-PPAR gamma (ab310323, 1:2,000, Abcam, Cambridge, MA, USA), anti-aggrecan (ab36861, 1:2,000, Abcam), anti-collagen II (ab188570, 1:2,000, Abcam), anti-MMP-13 (ab39012, 1:2,000, Abcam), anti-MMP3 (ab52915, 1:2,000, Abcam), anti-Beclin 1 (ab207612, 1:2,000, Abcam), anti-BCL-2 (ab182858, 1:2,000, Abcam), anti-Bax (ab32503, 1:2,000, Abcam), LC3A/B (4108, 1:1,000, Cell Signaling Technology, Danvers, MA, USA), anti-β-actin (ab6276, 1:2,000, Abcam), p44/42 MAPK (Erk1/2) (137F5) Rabbit mAb (4695, 1:1000, Cell Signaling Technology), Phospho-p44/42 MAPK (Erk1/2) (Thr202/Tyr204) (D13.14.4E) XP® Rabbit mAb (4370, 1:1000, Cell Signaling Technology), anti-CD9 (ab307085, 1:1000, Abcam), anti-CD63 (ab108950, 1:1000, Abcam), anti-ALIX (ab275377, 1:1000, Abcam), anti-Calnexin (ab22595, 1:1000, Abcam), anti-Hrs (sc-271455, 1:1000, Santa Cruz Biotechnology, Santa Cruz, CA, USA), STAM1 Antibody (13053, 1:1,000, Cell Signaling Technology, Danvers, MA, USA), anti-CHMP7 (sc-271805, 1:1000, Santa Cruz Biotechnology) and anti-TSG101 antibody (ab133586, 1:1000, Abcam).

### 4.11 Autophagy Flux Detection

Primary chondrocytes treated with FLS-derived exosomes were placed onto cell climbing slides and transfected with the mRFP-GFP-LC3 adenovirus (MOI = 10; polybrene = 2 μg/mL, provided by Hanbio), either before or simultaneously. Following the procedure outlined in Section 2.5, the chondrocytes were then treated and subsequently fixed in 4% paraformaldehyde for 30 min at room temperature. Imaging was carried out using a two-photon fluorescence microscope (Olympus).

### 4.12 Immunofluorescence Microscopy

FLSs were seeded onto coverslips in a 24-well plate. Once 80% confluency was reached, coverslips containing FLSs were fixed with 4% paraformaldehyde for 20 min and then permeabilized with 0.5% Triton X-100 (Solarbio Science & Technology Co., Ltd., Beijing, China) for 20 min at 27°C. Next, coverslips were blocked with 5% BSA for 30 min without washing, and then incubated with PPARγ antibody (ab310323, 1:500, Abcam) at 4°C overnight. After washing thrice in PBS, cells were incubated with anti-rabbit IgG (H + L) F (ab′)2 fragment (Alexa Fluor® 488 conjugate) (4412s, 1:200, Cell Signaling Technology) for 3 h at 25°C. Nuclei were stained with 4,6-diamidino-2-phenylindole (DAPI) for 5 min. The endosomal vesicle structures were stained with FM4-64 (HY-103466, 6μM, MedChemExpress) for 5 min. Coverslips were visualized under a confocal microscope (Olympus).

### 4.13 Flow Cytometry

Chondrocyte apoptosis was evaluated utilizing the FITC Annexin V Apoptosis Detection Kit I (556547, BD Biosciences, San Jose, CA, United States). Following three successive washes with PBS, cells presenting membrane pores were stained with propidium iodide (PI) (BD Biosciences) and quantified using a FACSCalibur flow cytometer (BD Biosciences). The rate of cell mortality was then normalized to that of the control group, and the resulting data were subsequently analyzed using the FlowJo software (v.10.7, BD Biosciences).

### 4.14 Transmission Electron Microscopy

In line with prior studies, the cells were first fixed in 2.5% glutaraldehyde for a period of 2-4 h at room temperature. After performing low-speed centrifugation at 500 xg for 5 min, cell clusters roughly the size of mung beans could be observed at the bottom of the tube. These clusters were then coated with 1% agarose and subsequently rinsed three times using a 0.1 M phosphoric acid buffer (PB, pH 7.4), with each rinse lasting for 15 min. Subsequently, 1% OSO4 was gently added to elevate and resuspend the cell clusters. After dehydration, the samples were embedded in resin. Following a uranium-lead double staining process (employing 2% uranium acetate saturated alcohol solution and lead citrate, each for a duration of 15 min), the sections were left to dry overnight at a temperature of 25°C. The cellular morphology and subcellular structures were then observed using a Hitachi 800 transmission electron microscope (TEM) (Tokyo, Japan).

### 4.15 Chromatin Immunoprecipitation-quantitative Polymerase Chain Reaction (ChIP-qPCR)

We employed ChIP-qPCR to examine the interaction between PPARγ and ANXA1. Cellular lysates from FLSs were harvested and a ChIP assay was performed using the SimpleChIP® Plus Enzymatic Chromatin IP Kit (9005, Cell Signaling Technology). In brief, FLSs were treated with 1% formaldehyde for 10 min to establish cross-links between DNA and protein. Fragmented chromatin was then extracted using 0.5 μL of micrococcal nuclease (10011; CST) and ChIP buffer. These were incubated overnight with rabbit IgG (2729; CST) as a control or anti-PPARγ antibody (sc-7273, Santa Cruz Biotechnology, Santa Cruz, CA, USA) at 4°C. The resulting mixtures of protein-DNA and antibody were further incubated with ChIP-Grade Protein G Magnetic Beads (9006, Cell Signaling Technology). After sequential washing with low- and high-salt solutions, the protein-DNA crosslinks in the immunoprecipitates were uncoupled using ChIP Elution Buffer (7009, Cell Signaling Technology). The DNA was subsequently purified using DNA purification columns, and this purified DNA was subjected to qPCR using ANXA1 primers supplied by Sangon (China; see supplementry table 1).

### 4.16 Co-Immunoprecipitation (Co-IP)

For Co-Immunoprecipitation (Co-IP), cells were lysed using IP/Co-IP lysis buffer (Cat# PC102, Epizyme, Shanghai, China). The lysates were subsequently immunoprecipitated using anti-PGC-1α antibody (D-5) (sc-518025, 1:30, Santa Cruz Biotechnology, CA, USA), anti-ANXA1-antibody (ab214486, 1:30, Abcam), p44/42 MAPK (Erk1/2) (137F5) Rabbit mAb (4695, 1:30, Cell Signaling Technology), anti-Hrs (sc-271455, 1:30, Santa Cruz Biotechnology), STAM1 Antibody (13053, 1:30, Cell Signaling Technology), anti-CHMP7 (sc-271805, 1:30, Santa Cruz Biotechnology) and anti-TSG101 antibody (ab133586, 1:40, Abcam). A total of 2μg rat IgG antibody (A7031, Beyotime, Shanghai, China) was used as an internal control. These components were mixed with Protein A/G magnetic beads (HY-K0202, MedChemExpress, Monmouth Junction, NJ, USA) and incubated at 4℃ with rotation overnight. Following incubation, the mixture was washed thrice with 1 mL of lysis buffer, and the protein complexes were eluted by boiling in SDS sample buffer. The resulting precipitated protein was then analyzed via Western blotting using the aforementioned antibodies.

### 4.17 Immunohistochemistry

Briefly, following the kit manufacturer’s instructions (CAT # SP-9001, Zhongshan Jinqiao, China), we performed immunostaining of proteins using a two-step method. The antibodies used in this part of the study were the same as those used in western blotting. Procedures were carried out as we have previously described [9, 10]. The average optical density was used to represent the relative expression of aggrecan and type II collagen, while the expression of MMP13, MMP3, Beclin-1, BCL-2, and Bax was quantified based on the percentage of cells that stained positive.

### 4.18 Statistical Analysis

Results are presented as means ± standard error of the mean (SEM), analyzed using GraphPad Prism 5 (GraphPad Software Inc, San Diego, California, USA). Statistical evaluations were conducted using Student’s t-test and one-way ANOVA, facilitated by IBM SPSS Statistics 25.0. A p-value of < 0.05 was considered statistical significant.

## Declarations

## Acknowledgements

We appreciate the help from colleagues in the affiliated Laboratory of China Medical University and Beijing Seekgene Biotechnology Co., LTD

## Author contributions

Conceptualization: Shuangshuo Jia, Lunhao Bai; Methodology: Shuangshuo Jia; Formal analysis and investigation: Jiabao Liu, Ziyuan Wang; Writing - original draft preparation: Shuangshuo Jia; Writing - review and editing: Shuangshuo Jia, Lingyu Yue, Yingliang Wei; Funding acquisition: Yue Yang, Lunhao Bai; Resources: Shuangshuo Jia, Lunhao Bai; Supervision: Lunhao Bai

## Ethics approval

The studies involving human participants were reviewed and approved by the protocol and experiments on joint specimen after human replacement were approved by the Ethics Committee of Shengjing Hospital of China Medical University (No: 2019PS629K). The patients/participants provided their written informed consent to participate in this study. The animal study was reviewed and approved by the Ethics Committee of China Medical University (no. 2017PS237K). Written informed consent was obtained from the individual(s) for the publication of any potentially identifiable images or data included in this article.

## Consent to participate

Informed consent was obtained from all individual participants included in the study.

## Data availability

The data displayed in Figs are openly available in mendeley at https://data.mendeley.com/datasets/fm7gwd4rct/1. Any additional information required to reanalyze the data reported in this paper are available from the corresponding author upon reasonable request

---

**Supplementary figure 1.**
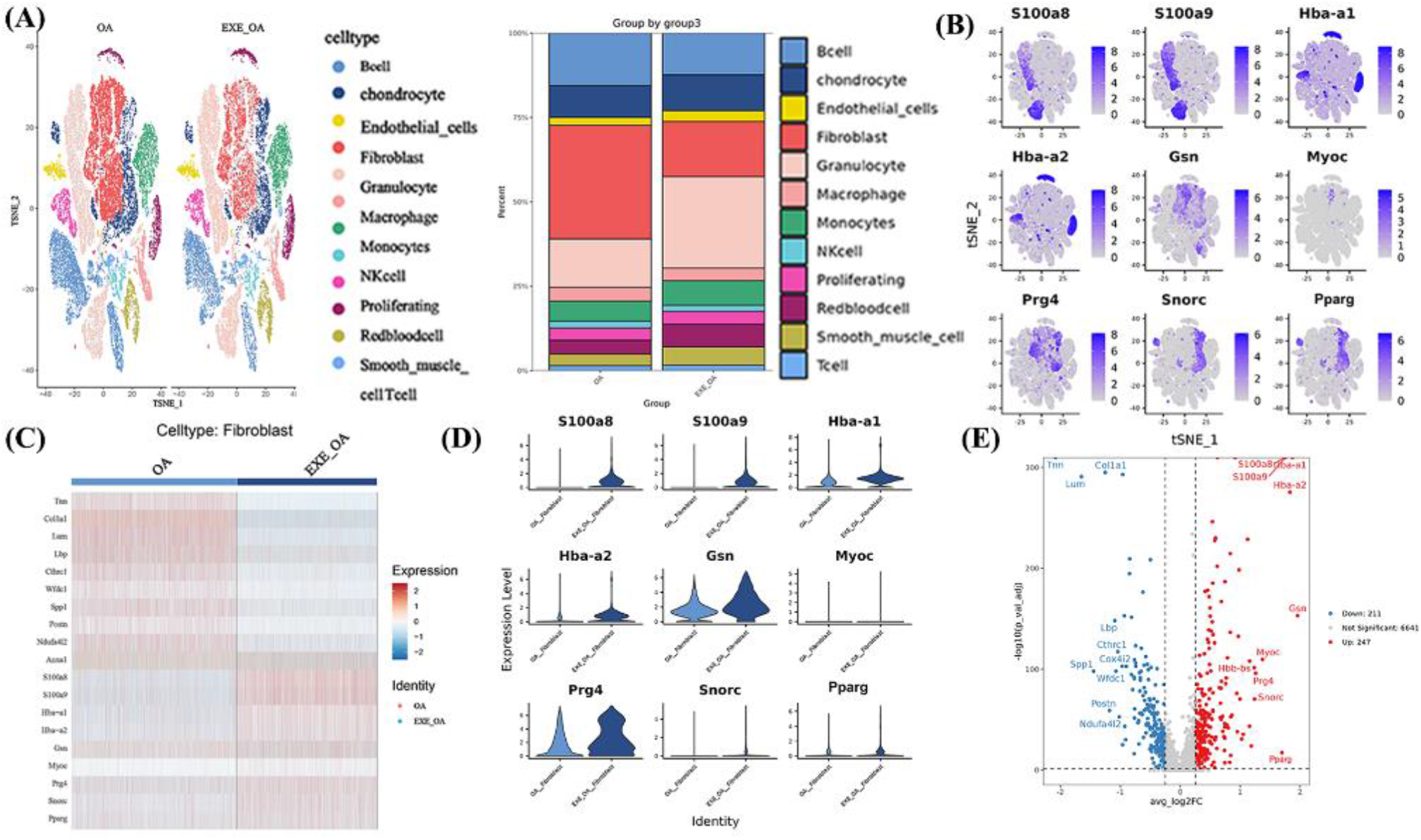
Increased expression of PPARγ in the FLSs of the moderate exercise group is implicated in OA pathogenesis, as determined by single-cell transcriptome sequencing. (A) A t-distributed stochastic neighbor embedding (t-SNE) map and an integrated percentage figure show the cell type distribution in the entire joint of SD rats. (B) The t-SNE map, (C) heatmap, (D) violin plot, and (E) volcano plot illustrate the top 9 upregulated genes in the FLSs when comparing the OA+exercise group versus the OA group. Among these genes, we found PPARγ to be upregulated in the FLSs of the OA+exercise group. (F) The ratio of relative protein expression in chondrocytes. In the 3-MA and Honokiol treatment series, *p < 0.05 vs. CG; **p < 0.01 vs. CG; ***p < 0.001 vs. CG. #p < 0.05 vs. IL-1β; ##p < 0.01 vs. IL-1β; ###p < 0.001 vs. IL-1β. +p < 0.05 vs. ANXA1; ++p < 0.01 vs. ANXA1; +++p < 0.001 vs. ANXA1. In the BOC-2 treatment series, *p < 0.05 vs. IL-1β; **p < 0.01 vs. IL-1β; ***p < 0.001 vs. IL-1β. #p < 0.05 vs. NC; ##p < 0.01 vs. NC; ###p < 0.001 vs. NC. +p < 0.05 vs. OE-PPARγ; ++p < 0.01 vs. OE-PPARγ; +++p < 0.001 vs. OE-PPARγ.

**Supplementary figure 2.**
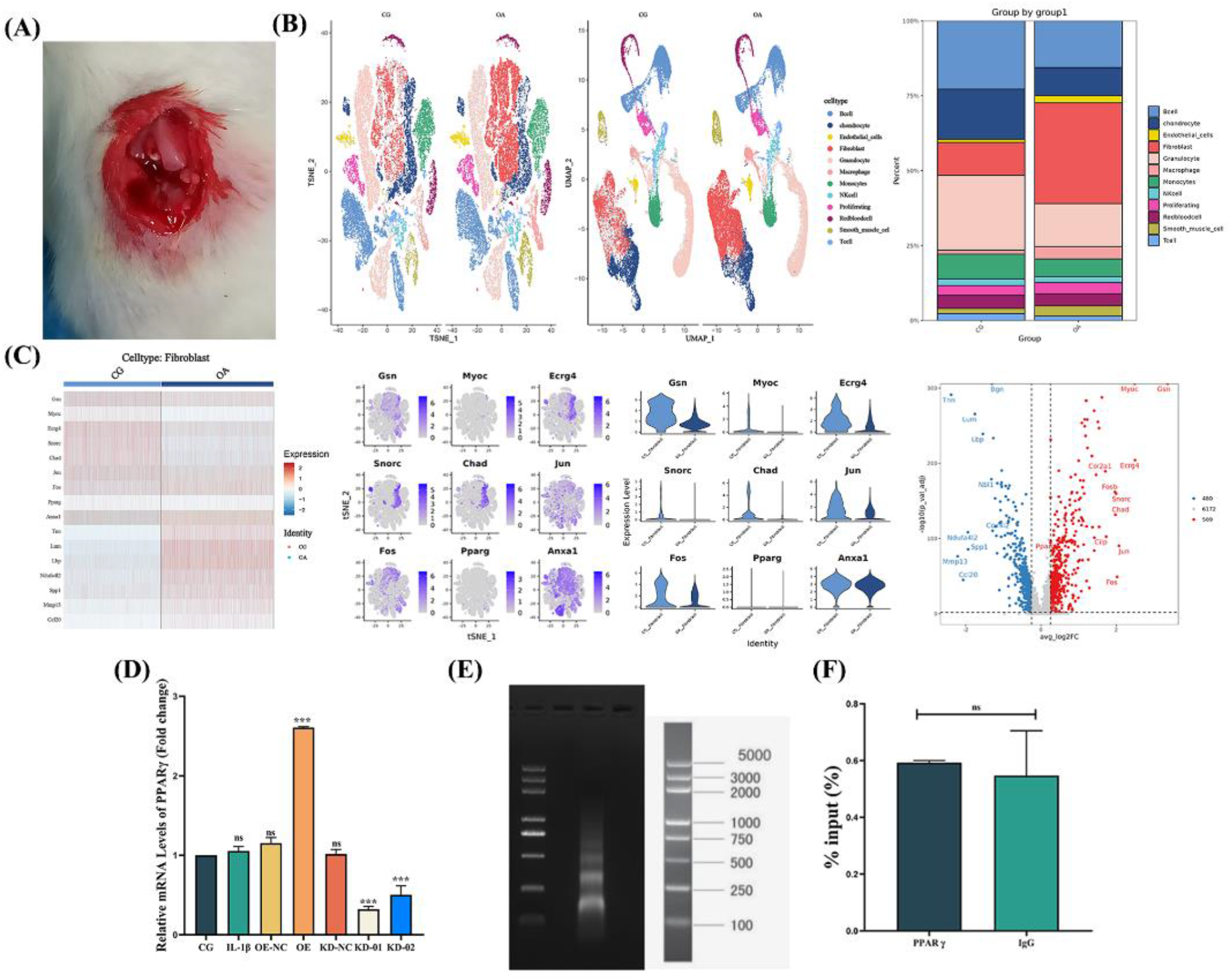
(A) The ACLT osteoarthritis model. (B) t-distributed stochastic neighbour embedding (t-SNE) and integrated percentage figure of the cell type of the whole joint of SD rats between CG and OA group. (C) The differentially expressed gene in FLSs between CG and OA group displayed in heatmap, t-SNE, violin plot and volcano plot. (D) The relative mRNA levels of PPARγ treated FLSs to evaluate the OE or KD efficiency. (E) Results of chromatin agarose gel electrophoresis in Chip-qPCR experiment. (F) Immunoprecipitated DNA using chromatin immunoprecipitation (ChIP) assay was quantified by qPCR in FLSs, with normal rabbit IgG as the negative control.

**Supplementary figure 3.**
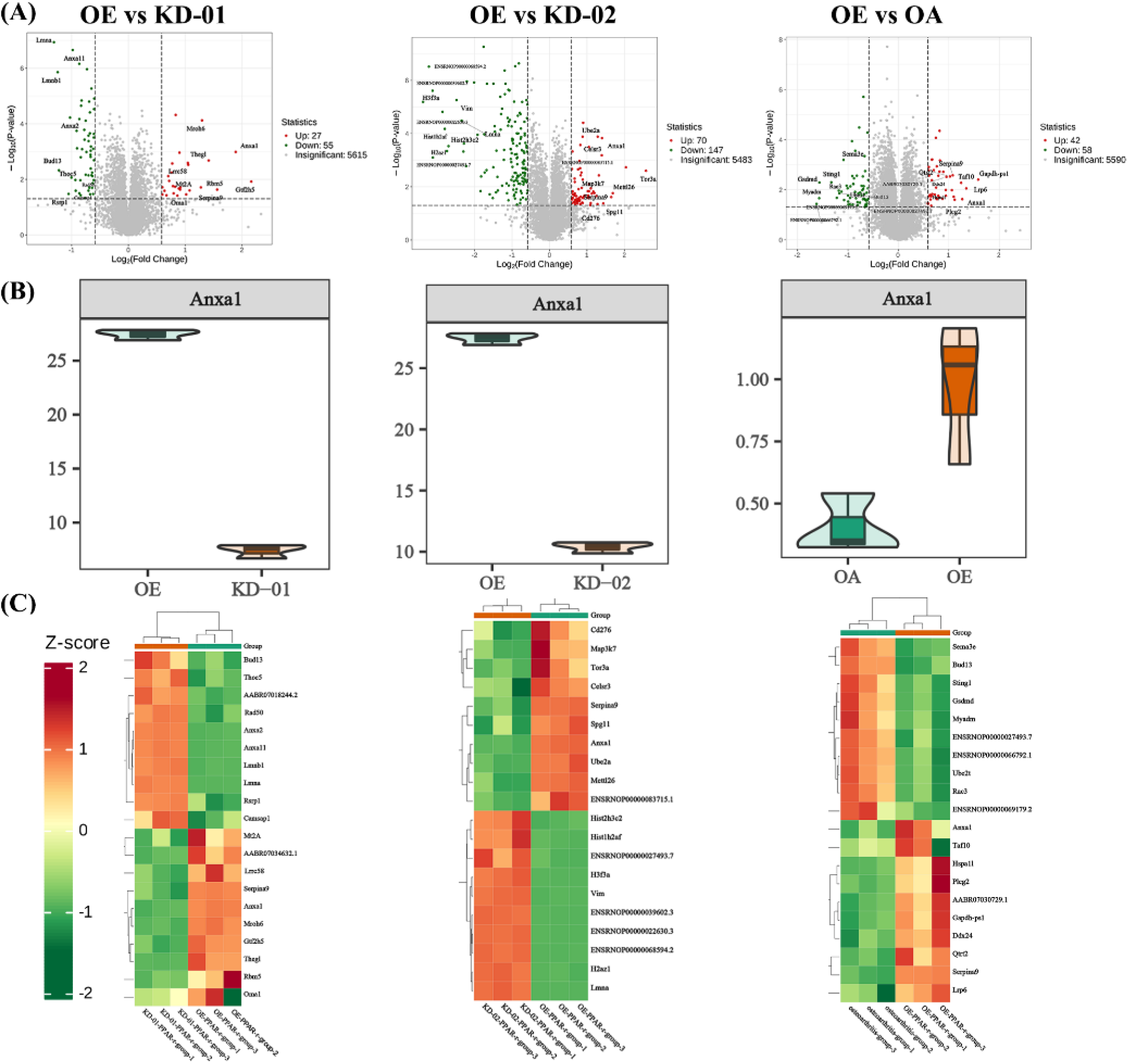
Annexin A1 is enriched in exosomes derived from OE-PPARγ FLSs. Using data-independent acquisition quantitative proteomics of FLS-derived exosomes, we found Annexin A1 to be enriched in exosomes derived from OE-PPARγ FLSs compared to KD-01, KD-02, and OA groups. The volcano plot (A), violin plot (B), and heatmap (C) depict the enrichment of ANXA1 in the OE-PPARγ group.

**Supplementary figure 4.**
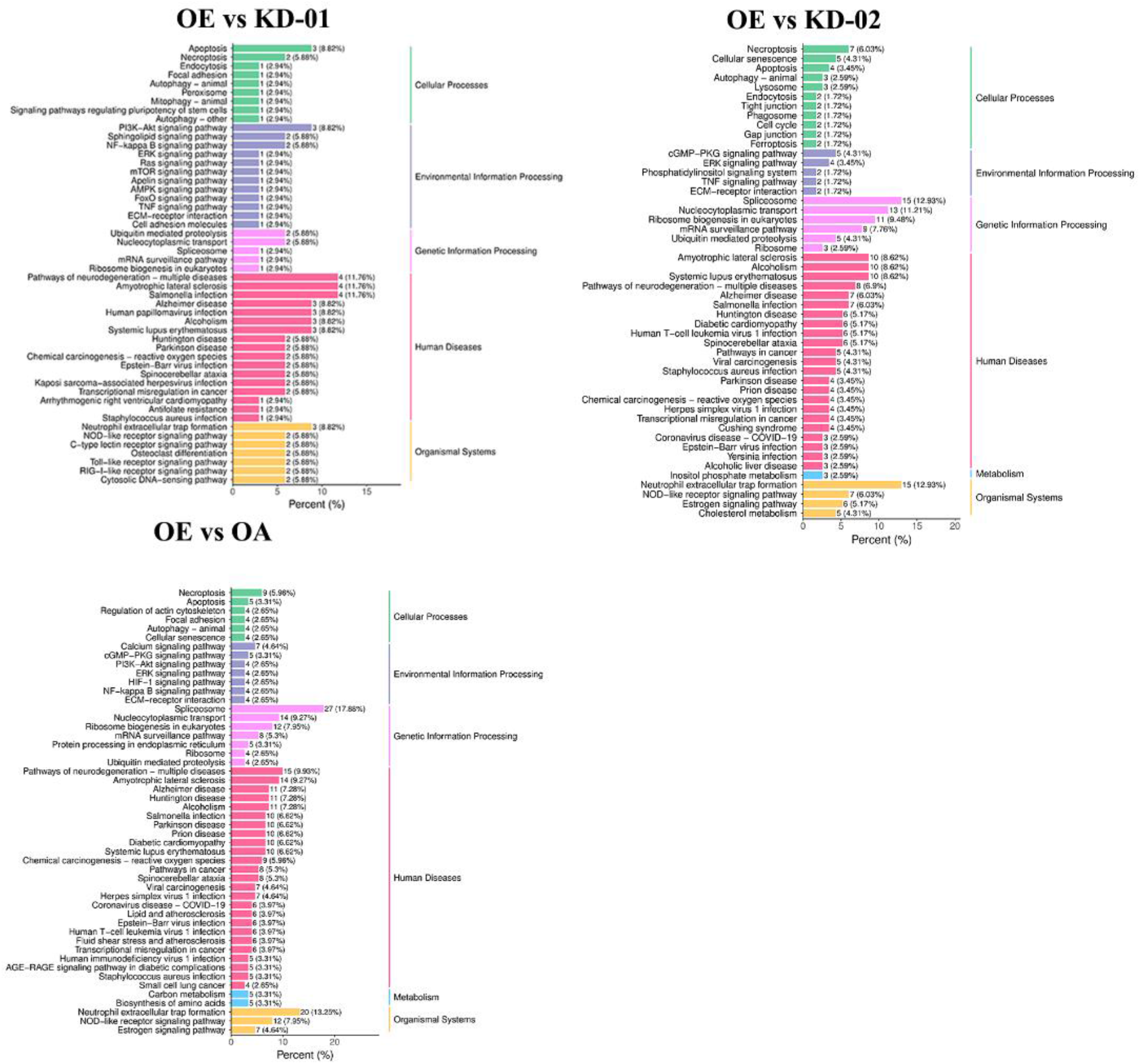
GO analysis of OE vs KD-01, KD-02, OA group.

**Supplementry table 1.**
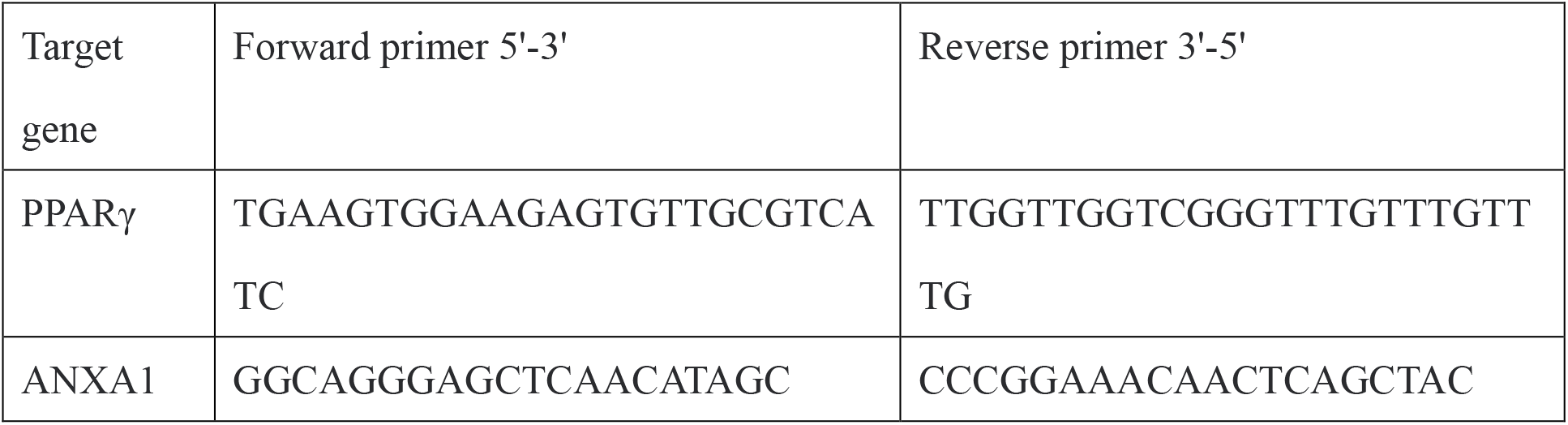
Sequences of primers used for qRT-PCR and Chip-qPCR.

